# A multi-ring shifter network computes head direction in zebrafish

**DOI:** 10.64898/2025.12.29.696831

**Authors:** Siyuan Mei, Hagar Lavian, You Kure Wu, Martin Stemmler, Ruben Portugues, Andreas V.M. Herz

**Affiliations:** Faculty of Biology, Ludwig-Maximilians-Universität München, Munich, Germany; Graduate School of Systemic Neurosciences, Ludwig-Maximilians-Universität München, Munich, Germany; Bernstein Center for Computational Neuroscience Munich, Munich, Germany; Institute of Neuroscience, Technical University of Munich, Munich, Germany; SyNergy Excellence Cluster, Munich, Germany; Max Planck Fellow Group - Mechanisms of Cognition, MPI Psychiatry, Munich, Germany; Department of Neurobiology, George S. Wise Faculty of Life Sciences, Tel-Aviv University, Tel-Aviv, Israel; Institute of Neuronal Cell Biology, School of Medicine, Technical University of Munich, Munich, Germany; Department of Neurobiology and Behavior, Cornell University, Ithaca, New York, 14853, USA

## Abstract

From insects to fish to mammals, many species have an internal compass: a set of recurrently connected neurons that combine motor feedback, vestibular signals, and external cues to compute the animal’s heading direction. Whether the underlying mechanism is universal across different species is unresolved. In Drosophila, for instance, the central complex contains three connected, yet anatomically separate, functional neuron rings. Two rings receive countervailing velocity signals that shift neuronal activity bumps around all three rings. An alternative, classical continuous-attractor model invokes only one ring but requires velocity-modulated synaptic connections. A single-ring compass has been discovered in the anterior hindbrain of zebrafish. Using theory and experiments, we find, however, that three rings, including two shifter rings, are intermingled on the same anatomical scaffold. Zebrafish and Drosophila, therefore, use the same basic compass mechanism, suggestive of convergent evolution.

## Introduction

Even in darkness, many animals retain a sense of direction because their nervous system keeps a memory of heading ^1,2^; moreover, they update this memory by integrating angular head velocity (AHV) as relayed by the vestibular system ^3^, optic flow, or motor efference signals ^4^. Other sensory cues, such as landmark information ^5^, are used to calibrate and correct this error-prone integrating mechanism ^2,6^. At the neuronal level, when an animal faces a given azimuthal direction, a particular set of head-direction (HD) cells fires ^1–6^. When the animal rotates, the set of active HD cells is updated such that, as a whole, the HD cell population acts as an internal compass.

Two alternative mechanisms for HD networks have been put forward. Skaggs et al. ^7^ proposed that HD cells form a (functional) ring of neurons whose population vector signals HD. A turn by the animal activates cells that rotate the localized bump of the HD cell population. These “rotation” or “shifter” cells form two additional rings, one sensitive to clockwise (CW) rotations and one for counterclockwise (CCW) rotations. This is consistent with Drosophila’s HD system, which has separate anatomical structures that instantiate the three rings ^8–11^. The second mechanism eliminates shifter cells entirely ^12^ and assumes that AHV-modulated synaptic couplings move the activity bump around a single ring.

HD cells have been identified in many species, including Drosophila ^2^, desert locusts ^13^, mice ^3^, rats ^5^, bats ^14^, and fish ^15,16^. This raises the question of which mechanism different species use to rotate the HD representation.

The tuning properties of those HD cells that are also sensitive to AHV might provide an answer. For the synapse-modulation mechanism proposed by Zhang ^12^, neurons are only tuned to HD. In models with shifter cells, however, conjunctive HD × AHV tuning is an *obligatory consequence*: indeed, for the shifter mechanism to function properly, the activity difference between the two shifter rings must depend on AHV. This mathematical requirement results in asymmetric AHV tuning of the shifter rings such that the activity skews either to the CW or CCW direction.

Here, we present a framework to distinguish these alternative possibilities using neuronal activity alone. We discuss the HD systems in Drosophila and zebrafish and show that, in both species, HD cells have V-shaped AHV tuning, meaning that the activity of these cells is minimal when the animal does not rotate and increases for both positive and negative AHVs. For some cells, these two slopes differ, resulting in skewed AHV tuning, whereas for other cells, AHV tuning has no skew. Classifying Drosophila HD cells based on their AHV skew yields a tight match with the three known anatomical rings. When we apply this same analysis to zebrafish, we find a balanced number of CW-and CCW-shifter cells. Anatomically, the shifter rings are partially lateralized, and their preferred HDs are non-uniformly distributed across all 360°. We develop a ring-attractor network model that accounts for such non-uniformity. Our theoretical framework can be readily applied to other species for which the anatomical layout of the HD system is unknown.

## Results

### Two alternative compass mechanisms

Across species, the neuronal representation of the 360-degree heading is hypothesized to reside in ring-attractor networks ^7,12,17^. In these models, the synaptic strength between two cells depends on the angular distance between their preferred HDs, yielding a synaptic coupling matrix that is symmetric and rotation invariant. Typically, local neighbors, measured in functional space, are connected by excitatory couplings, while neurons that are further away inhibit each other. Figure 1a depicts the essential wiring characteristic (left panel) and an example of synaptic connectivity (right panel). For sufficiently strong couplings, the network evolves towards a stationary bump of activity on the ring (Fig. 1a, left panel). To serve as a compass, an update mechanism is needed to integrate AHV signals when the animal turns to yield its HD, moment by moment.

**Fig. 1:**
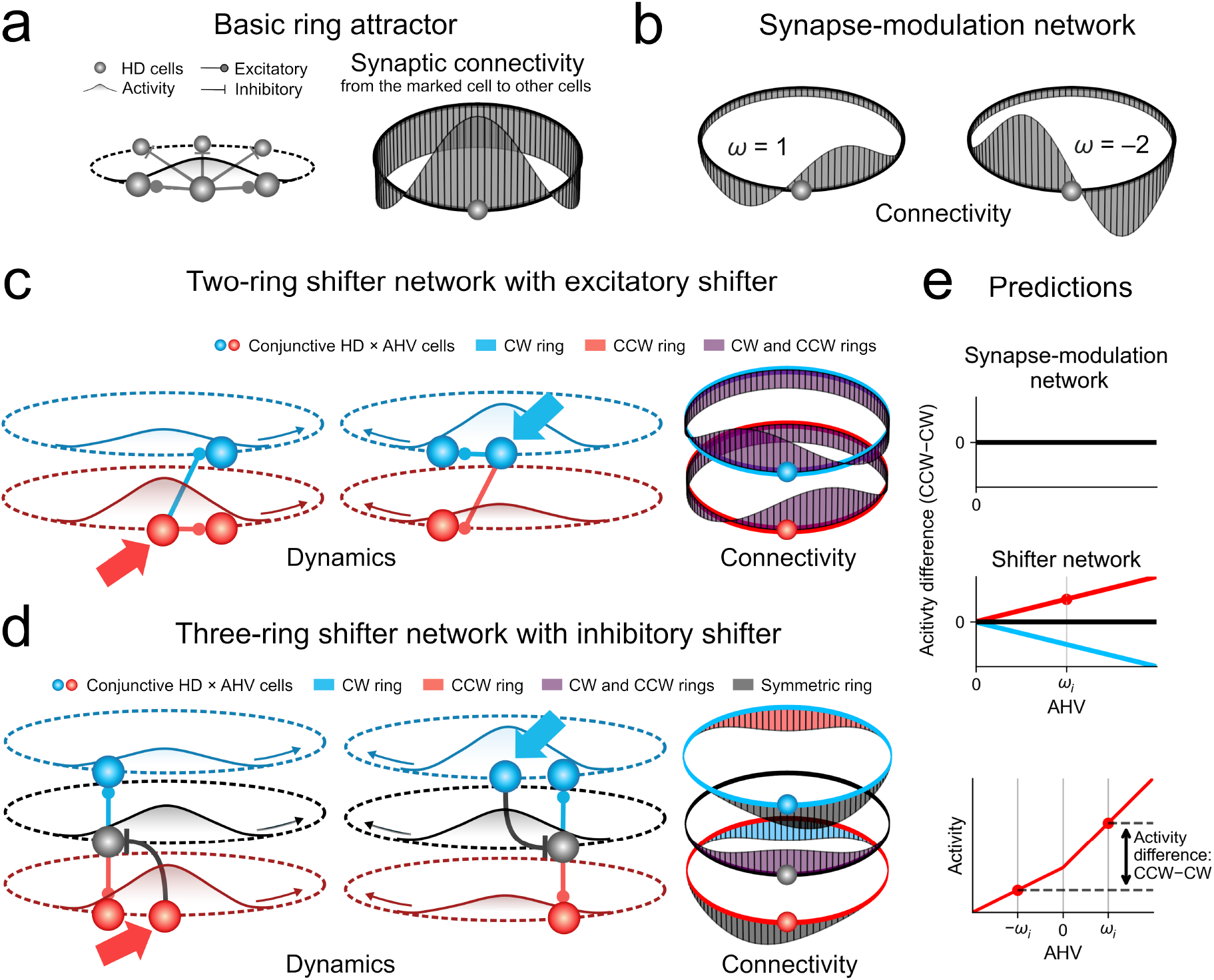
Tuning to angular head velocity distinguishes two classes of ring-attractor networks. **a**, Arranging head-direction (HD) cells as a ring and using locally excitatory and long-range inhibitory synaptic connections creates a stationary attractor that maintains directional information but cannot integrate angular head velocity (AHV) to yield HD. **b**, Networks with AHV-dependent modulation of synapse-strength asymmetry can integrate AHV. When an animal rotates counterclockwise (CCW, left panel), the synaptic strength is skewed CCW and scaled by the angular-head speed (AHS), driving the population activity CCW. The same is true for clockwise rotations (CW, right panel). **c**, Alternatively, in shifter networks, countervailing velocity inputs to a pair of neuron rings underlie AHV integration. When an animal rotates CCW, the CCW ring receives stronger signals (large red arrow) and becomes more active. In combination with the CCW-skewed excitatory connections, the population activity is shifted CCW (left panel)—and vice versa for CW rotations (right panel). **d**, A variant of (c) with an additional symmetric (HD readout) ring projecting to both shifter rings. Shown is an example with inhibitory connections from the CCW (CW) ring that are skewed CW (CCW); here, the symmetric ring excites both shifter rings without a skew. When an animal rotates CCW, the stronger activity of the CCW ring inhibits movement in the CW direction so that the population activity rotates in the CCW direction (left panel). The same is true for CW rotations (right panel). **e**, The activity difference between opposite rotations distinguishes synapse-modulation networks from shifter networks. The activity of synapse-modulation does not depend on the sign of AHV. In a shifter network, however, the activity difference (CCW-CW) increases for the CCW ring with increasing AHV, while the CW ring prefers negative AHV. Only the activity of the symmetric (HD readout) ring is symmetric, as in the synapse-modulation network.

Two mechanisms have been proposed that can move the attractor state to reflect the current HD. In the first category, the synapse-modulation network, AHV modulates the synaptic strength to make all couplings slightly asymmetric, but still rotation-invariant across the units in the network ^12,18,19^ (Fig. 1b; Methods, Eqs. (1) and (2)). This type of network has been suggested to account for the HD system in desert locusts ^20^.

The second mechanism invokes explicit “rotation” or “shifter” cells (Methods, Eqs. (3) to (8)), which are arranged on two separate rings that project to a central ring of HD neurons, as proposed by Skaggs et al ^7^. Xie et al. ^21^ reduced such networks to just the two shifter rings (Fig. 1c) with no central ring at all. Neurons connect across rings (and, in some cases, also within rings) with a positional offset. When the animal turns counterclockwise, i.e., with positive AHV *ω*, all neurons in the CCW ring receive a biased input b(*ω*), while the neurons in the CW ring receive a weaker bias b(-*ω*), and vice versa for negative AHV. The positional shifts in the synaptic wiring will then cause activity on the rings to move in response to the bias difference. The same mechanism works when one reintroduces a central HD ring ^22^ (Figure 1d). Such a three-ring structure has been discovered in Drosophila ^9,10^.

In shifter models, neurons belonging to the CW and CCW shifter rings develop conjunctive HD×AHV tuning as an emergent property of the network dynamics (see Methods and Supplementary Information for analytical and simulation results). The AHV affects the relative amplitude of activity. In fact, we show that the activity difference between opposite rotations is approximately linearly proportional to AHV (Fig. 1e, Methods). Notably, the activity in both rings could still be at its lowest point when the velocity is zero, as long as the velocity-induced increase skews towards one direction. Crucially, this allows us to relax the usual assumption that b(*ω*) is a linear function of *ω*. Instead, we may write *b*_shift_ (*ω*) *g*(|*ω*|) + *kω*, where *g* can be any real function, |*ω*| is the absolute value of *ω*, and *k* is a positive number. This more flexible framework raises the prospect of identifying shifter circuits even when the synaptic wiring diagram is unknown, and the AHV-tuning is V-shaped. Activity levels in the synapse-modulation network proposed by Zhang ^12^, by contrast, will be invariant under variations of the AHV. This difference allows us to discriminate between the two mechanisms.

To establish a ground truth case for comparison, we first reanalyze Drosophila calcium imaging data. We then turn to HD tuning in zebrafish, for which the synaptic connectome is not fully known.

### Drosophila

The Drosophila HD system is thoroughly studied and described functionally and anatomically ^2,8–11^, making it the ideal system to test our theoretical framework. The fly’s central complex (Fig. 2a) contains three anatomically separate “rings” (Fig. 2b): EPG neurons (symmetric ring; pure HD cells), and PEN neurons (conjunctive HD × AHV cells), which are located on the fly’s left (PEN-L neurons) and right hemispheres (PEN-R neurons). The three rings constitute a shifter network (Fig. 2c). When the fly rotates CCW, left PEN neurons (PEN-L) receive stronger inputs, resulting in higher activity. Because the excitatory connections from PEN-L neurons to EPG neurons are skewed towards the CW direction, and EPG neurons excite both types of PEN neurons without a skew, the heightened PEN-L activity moves the population activity of all three rings in the CW direction. Likewise, when the fly rotates CW, PEN-R neurons increase their activity and rotate the bump counterclockwise. Unlike PEN-R or PEN-L neurons, cells in the symmetric ring (EPG) show no preference for rotation direction (Fig. 2d, left panel).

**Fig. 2:**
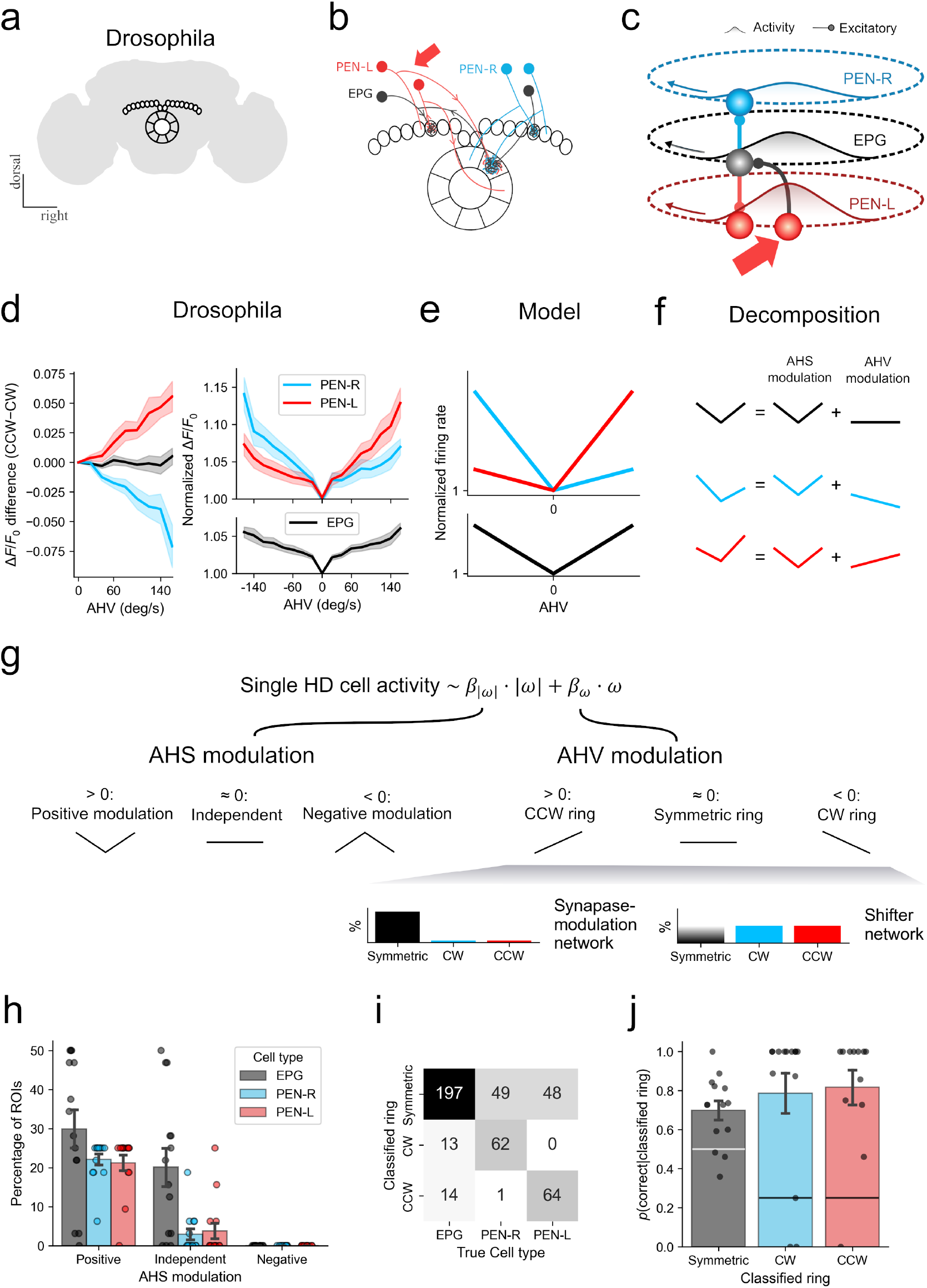
The Drosophila head-direction system is consistent with a shifter network but not a synapse-modulation mechanism. **a**, The fly’s HD system is located in the central complex. **b**, The core HD system consists of EPG neurons (pure HD tuning) and left and right PEN1 neurons (PEN-L, PEN-R, respectively; conjunctive tuning to angular head velocity and HD, i.e., AHV-HD tuning). **c**, The HD system forms a three-ring shifter network. Connections from PEN-L (PEN-R) to EPG neurons are excitatory and skewed CW (CCW), EPG neurons excite PEN-L and PEN-R neurons without skew. When a fly rotates CCW (CW), PEN-L (PEN-R) neurons receive larger AHV signals (large red arrow) and fire more strongly, shifting the population activity in the opposite direction, i.e., CW (CCW). Compared to the conventional three-ring shifter network, this opposite rotation direction reflects the fly’s flipped anatomy but does not affect the rings’ overall function. **d**, The activity of all HD cells depends in a V-shaped manner on AHV, i.e., the cells are positively modulated by angular head speed (AHS), the absolute value of AHV (left panel). In PEN-L neurons, the activity difference for opposite rotations increases with AHV, while in PEN-R neurons, it becomes more negative. The activity of the symmetric ring exhibits no direction preference (right panel). **e**, In shifter networks, a positive AHS modulation may arise from increasing the external inputs with AHS. **f**, Neuronal activity levels can be split into two independent components: AHS modulation and AHV modulation. Three examples are shown. **g**, Linear regression reveals the contributions from AHS and AHV modulation. In addition to other predictors specific to the experiments, the activity of HD cells can be predicted by AHS (|*ω*|) and AHV (*ω*). A significantly positive (negative) AHS coefficient *β*_|*ω*|_ indicates a positive (negative) AHS modulation and a (inverted) V-shaped activity dependence, while a non-significant coefficient suggests that there is no AHS modulation. A significantly positive (negative) AHV coefficient *β*_*ω*_ points to a CCW (CW) rotation preference, i.e., a CCW (CW) ring, while a non-significant coefficient suggests no direction preference, as expected for a symmetric ring. Near-zero *β*_*ω*_-values of almost all HD cells indicate a synapse-modulation mechanism. Otherwise, if the numbers of positive and negative *β*_*ω*_-values are (approximately) equal, this suggests a shifter network, regardless of the number of non-significant *β*_*ω*_-values. **h**, The fraction of each tuning category among all ROIs. Dots represent individual flies and bars show the average percentage across flies. **i**, Confusion matrix between the classified rings and measured cell types. **j**, The conditional probability, i.e., the fraction of correct classification given the classified ring, is calculated separately for each ring. Dots represent individual flies and the horizontal lines mark the null hypothesis.

Although angular head speed (AHS), i.e., the absolute value of angular head velocity, positively modulates the activity of all rings (Fig. 2d, right panel), this is still consistent with the shifter model because the speed modulation can be incorporated into the model by increasing the overall inputs for increasing AHS (Fig. 2e, Methods). The relation between AHV and the activity difference of opposite rotations remains approximately linear, as shown by extensive simulations (Supplementary Information) covering a large range of model parameters (mean Pearson’s r = 0.998, SD = 0.013). In summary, the response characteristics of each HD cell can be viewed as a sum of two independent components: AHS modulation and AHV modulation (Fig. 2f), analogous to the possibility of decomposing a function into a symmetric and an antisymmetric part. Individual HD cell responses can therefore be decomposed using linear regression as:

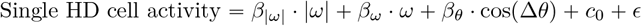

Here, *β*_|*ω*|_ captures the AHS modulation such that *β*_|*ω*|_ values significantly higher (lower) than zero correspond to a positive (negative) AHS correlation. *β*_*ω*_ captures the AHV modulation and thus includes the preference for the rotation’s direction. A cell with a significantly positive (negative) *β*_*ω*_ suggests a CCW (CW) rotation preference and therefore corresponds to the CCW (CW) ring, while a non-significant *β*_*ω*_ indicates no preference, as expected for the symmetric ring. If the *β*_*ω*_ values of almost all HD cells are non-significant, this points to a synapse-modulation mechanism. Otherwise, if equal numbers of significantly positive and negative *β*_*ω*_s are present, this suggests a shifter network, regardless of the number of non-significant *β*_*ω*_ values (Fig. 2g). Finally, Δ*θ* denotes the difference between the HD estimated from all HD cells, which corresponds to the current heading estimate, and the individual cell’s preferred HD. The term *c*_0_ represents baseline activity, and *ϵ* accounts for noise contributions.

We started by assessing the AHS modulation and the accuracy in assigning individual measurements to the three possible rings based on AHV modulation. We applied the above linear regression model to the calcium activity from the protocerebral bridge of 14 flies ^10^, with 16 EPG regions of interest (ROIs), eight PEN-L, and eight PEN-R ROIs in each fly. This revealed that 59.8% of EPG ROIs, 84.8% of PEN-L ROIs, and 88.4% of PEN-R ROIs were positively modulated by AHS (Fig. 2h). The classification performance was assessed using a confusion matrix (Fig. 2i) and conditional probabilities of correct assignment (Fig. 2j). These probabilities were above chance for both the shifter rings (CW: 0.787, Wilcoxon signed-rank test: *W* ^+^ = 88.0, *N* = 13, *p* = 1.01 × 10^−3^; CCW: 0.816, *W* ^+^ = 76, *N* = 12, *p* = 7.32 × 10^−4^; chance: 0.25) and the symmetric central ring (0.700, *W* ^+^ = 97.0, *N* = 14, *p* = 1.52 × 10^−3^; chance: 0.5). The tight match between the neurons’ assignment and their true identity demonstrates the general applicability of our theoretical framework to data from HD-cell networks.

### Zebrafish

HD cells have been identified in zebrafish ^16^. Their cell bodies are located in the anterior hindbrain, and their axons and dendrites project into the interpeduncular nucleus (Fig. 3a,b). The preferred HDs of these cells show some partial topographical organization (Fig. 3c), with inhibitory connections occurring between cells that are approximately 180 degrees apart in terms of their preferred HDs (Fig. 3d). Although the network updates its heading when the fish turns, the mechanism by which AHV is integrated remains unknown.

**Fig. 3:**
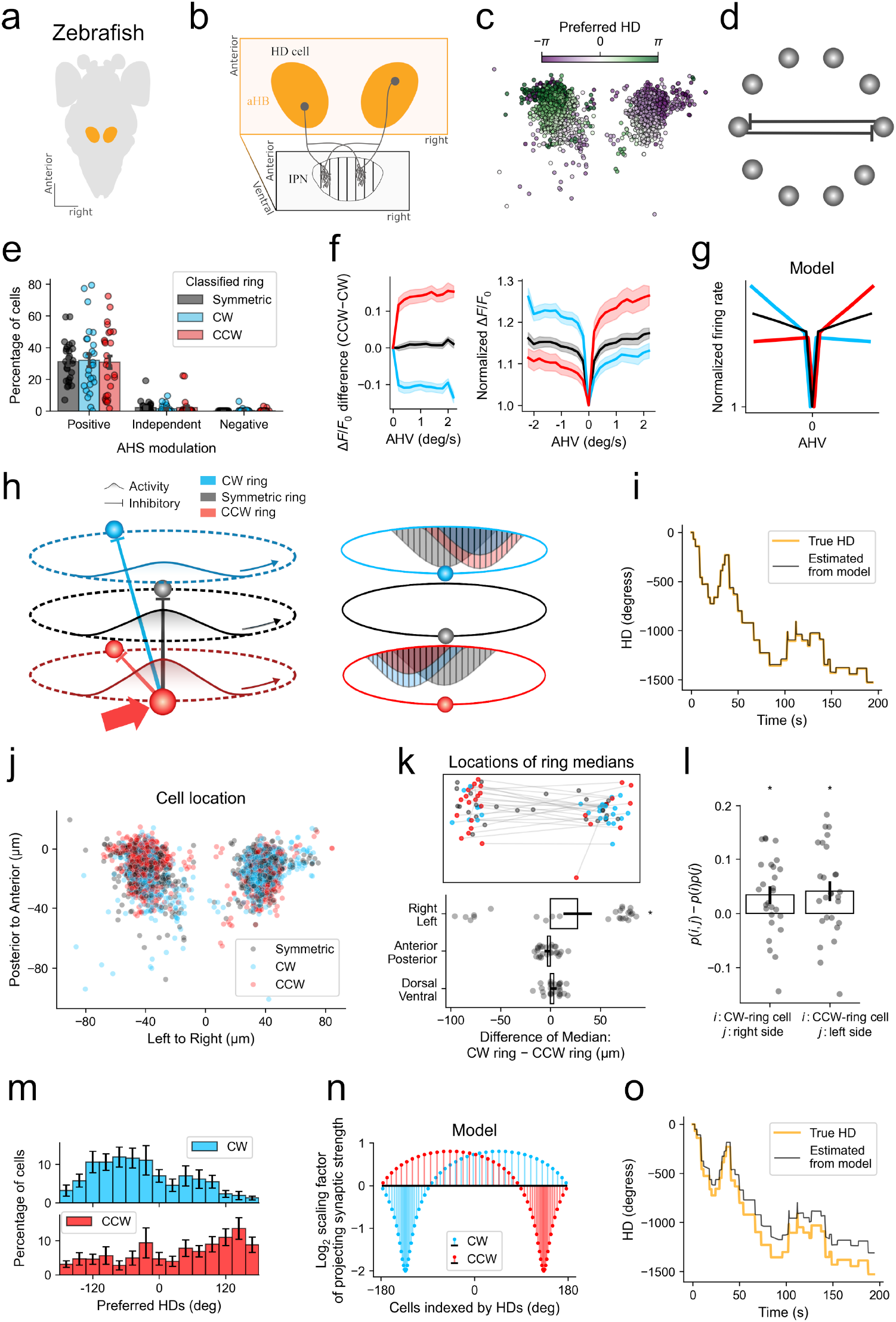
The zebrafish head-direction system is consistent with a shifter network but not a synapse-modulation mechanism. **a**, Cell bodies are located in the anterior hindbrain (aHB). **b**, Cell axons and dendrites synapse in the interpeduncular nucleus (IPN). **c**, Preferred HDs are approximately topographically organized. **d**, Connectome data suggest inhibitory connections to occur among cells whose preferred HD is roughly 180 degrees apart. **e**, Linear regression reveals that the number of symmetric, CW, and CCW cells is roughly the same, consistent with a shifter network. In addition, almost all HD cells are positively modulated by AHS. **f**, The CCW (CW) ring prefers CCW (CW) rotations, and the activity of the symmetric ring shows no preference. When the AHV changes from zero to non-zero levels, the activity of all neurons first increases abruptly, without skew for the symmetric ring and asymmetrically for the CW and CCW ring. To isolate the AHV modulation, the variance attributable to HD preference was removed based on linear regression. **g**, The strong increase can be incorporated into a ring-attractor model by a corresponding increase of the inputs. **h**, Sketch of the zebrafish HD shifter-network model. All neurons are inhibitory and receive a global excitatory AHV input. The left panel shows the basic network motif: Inhibitory connections from the CCW (CW) ring to both shifter rings are skewed CW (CCW). When zebrafish rotate CCW (CW), the CCW (CW) ring receives larger inputs and pushes the activity CCW (CW). The symmetric ring acts as a readout since the two shifter rings inhibit its most dissimilar HD representation. The right panel shows the synaptic connectivity from the cells marked for each ring to other cells in the same or other rings. **i**, Example for time-dependent HDs estimated from the zebrafish shifter network model using HDs and AHVs calculated from measured zebrafish trajectories. **j**, Anatomical locations of HD cells belonging to the CW, CCW, and symmetric rings for all fish. **k**, Upper panel: Median location of each ring of each fish. Lower panel: Distance between the median locations of the CW and CCW rings along the right-left, anterior-posterior, and dorsal-ventral axis. **l**, Joint probability of finding a CCW-ring cell on the left side, or a CW-ring cell on the right side, relative to the chance level, i.e., the product of the marginal probabilities. **m**, The distribution of the preferred HDs of the CW or CCW ring cells in zebrafish is non-uniform. **n**, For a ring-attractor model with non-uniform HD distributions, the projecting synaptic strength is scaled by the inverse density of preferred HDs. (**o)** As in (i) but using the model shown in (n).

To investigate the AHV integration and potential AHS modulations, we analyzed the calcium activity of HD cells in 27 zebrafish and applied the same linear regression as in Drosophila. The results revealed approximately equal proportions of HD cells belonging to the CW (0.337 ± 0.202) and CCW (0.331 ± 0.197) rings (paired-t test: *t* = 0.084, *p* = 0.934), indicating a shifter mechanism. AHS positively modulated the activities of most (2053 out of 2194) HD cells (Fig. 3e).

We next studied how the activity of each ring depended on AHV. As predicted, the CCW (CW) ring preferred positive (negative) AHV, whilst the symmetric ring showed no preference (Fig. 3f, right panel). Unexpectedly, the association between the activity difference of opposite rotations and AHV was not linear, possibly reflecting the influence of other neurons (bleed-through) on the imaging data or non-linearities in the calcium signal (Fig. 3f, left panel). In addition, the activity of all HD cells underwent an abrupt jump as AHV increased from zero to non-zero values (Fig. 3f, right panel, and Extended Data Fig.1). This feature can be incorporated into a shifter network by introducing a corresponding increase in the inputs driving the network (Fig. 3g).

To demonstrate the feasibility of our framework, we built a shifter network of the zebrafish HD system (see Methods, eqs. (15) –(17) and Supplementary Information), resembling the model of Boucheny et al. ^23^. To account for the known features of zebrafish HD cells, all neurons are inhibitory and receive a global excitatory input. The inhibitory connections from the CCW ring to the CW and CCW rings are skewed towards the CW direction; therefore, when zebrafish rotate CCW, the CCW ring receives higher inputs and pushes the activity to move in the CCW direction (and vice versa for CW rotations). The symmetric ring receives projections from the CW and CCW rings that inhibit the most dissimilar HD representation so that this ring can act as a readout. Given the morphology of the HD cells in zebrafish (Fig. 3b), we predict that the strongest connections within the shifter rings occur between cells with their preferred HDs slightly less than 180 degrees apart. In comparison, the strongest connections to the symmetric ring occur between cells exactly 180 degrees away (Fig. 3h). The resulting network accurately tracks HDs of zebrafish trajectories based on AHV inputs (Fig. 3i); the correlation between the HD estimated by network simulations and the true HDs approaches one (mean Pearson’s r = 0.995, SD = 0.048).

Inspired by the anatomical separation of shifter rings in Drosophila, we investigated the anatomical locations of the three rings in zebrafish. Cells belonging to the CW-ring tended to be located on the right side, while cells of the CCW-ring were predominantly present on the left side (Fig. 3j and Extended Data Fig.2). The median location of the CW ring was more right than the CCW ring (paired sample t-test: t(26) = 2.102, p = 0.046). The median location of the CCW and CW rings was not different along the anterior-posterior (t(26) = –1.608, p = 0.120) or dorsal-ventral axis (t(26) = 1.963, p = 0.061). The probability of finding a CW-ring cell on the right side or a CCW-ring cell on the left side was above chance (Wilcoxon signed-rank test: *W* = 92, *N* = 26, *p* = 0.019; *W* = 97, *N* = 26, *p* = 0.013, respectively) (Fig. 3l).

Given the topographic organization of preferred HDs and the anatomical lateralization of the CW and CCW rings, we expect a non-uniform distribution of preferred HDs within the CW and CCW rings. Indeed, neurons of the CW-ring preferentially encode HDs between -180° and 0° (The non-uniformity of 10 individual fish reached significance in the Rayleigh test after Bonferroni correction. Combining p-values using Pearson’s method renders a population level *p* = 1.21 × 19^-15^), while CCW-ring cells preferentially encode HDs between 0° and +180° (15 fish exhibited significance, population level *p* = 3.03 × 10^-17^) (Fig. 3m and Extended Data Figs. 3a and 3b). This non-uniformity poses a challenge for ring-attractor networks, since it destroys the rotational symmetry and reduces a continuous ring attractor to a small number of stable fixed points. However, this potential shortcoming can be compensated for by scaling the synaptic strengths by the inverse density of preferred HDs (Fig. 3n). The resulting network still tracks HDs, although its performance is slightly lower (mean Pearson’s r = 0.981, SD = 0.113) (Fig. 3o).

## Discussion

Head-direction (HD) cells provide a stable neuronal representation of an animal’s orientation in space and enable precise navigation, as shown by ablation experiments ^24^. Experimental evidence suggests that HD cells are arranged as one or multiple (functional) rings and establish a continuous attractor network. HD-cell activity forms a localized bump that stays still when the animal is at rest. When the animal rotates, the bump moves, serving as an internal compass that supports path integration.

Two mechanisms have been proposed to explain the compass rotation. In networks with synapse modulation, synapses within a single ring are modulated by angular head velocity (AHV); this results in asymmetric connections that push the activity bump around the ring ^12^. In shifter networks, on the other hand, two rings are tuned for clockwise and counterclockwise rotations of the animal, respectively ^7^. In some models, a third ring is added that is AHV-insensitive and serves as HD readout. Our mathematical framework shows that for a large class of shifter networks, the activity difference between opposite rotations is approximately proportional to the animal’s AHV. According to the synapse-modulation mechanism, however, the bump activity does not change at all with AHV ^12^. Together, these characteristic features allow us to use linear regression to discriminate between the synapse-modulation and shifter mechanisms, and to assign individual HD cells to their respective rings. This ability to infer network structure from neuronal function alone is crucial for understanding the HD mechanism in animal models, in which the connectome is unknown.

When we applied our approach to Drosophila, for which the ground truth is known, we recovered the shifter mechanism discovered in earlier studies ^9,10^. These results were made possible by the unique advantages that Drosophila offers: a small brain, individually identifiable neurons, a broad array of transgenic tools, and access to ultrastructural data. Applied to zebrafish ^16^, our analysis suggests a shifter mechanism similar to Drosophila, with three functional rings that are partially segregated anatomically. According to the experimental data, unlike in the fly, all three rings consist of inhibitory HD cells, which receive global excitatory input from a yet unknown source. Our findings suggest that in Drosophila and zebrafish, a three-ring compass structure serves as a basic computational motif, whose detailed neuronal composition is species-dependent.

Based on our analysis, we developed a ring-attractor model for zebrafish and predicted that the strongest connections among neurons in the shifter rings occur between cells with preferred HDs slightly less than 180° apart, as observed experimentally. We also found that the distribution of preferred HDs is non-uniform. This observation violates the rotational symmetry assumed in previous ring-attractor network models. But as we could also show, the continuous-attractor properties can be reestablished once the synaptic strength from HD cells in the shifter rings is inversely proportional to the HD density.

In zebrafish, nearby non-HD neurons show similar AHV preferences and partial lateralization (see Supplementary Information, Figs. S1 and S2), which possibly reflect common motor inputs to this brain region. Alternatively, this observation may indicate that the three-ring prediction is influenced by experimental signal bleed-through from neighboring non-HD neurons. Another potential explanation is that the data was collected from zebrafish larvae, and the system may change as the animal develops into the adult form.

The abrupt and strongly non-linear dependence of the activity difference on AHV in the zebrafish HD system violates the prediction of shifter networks. However, there are at least five possible explanations for this phenomenon. First, by experimental design, all recorded neurons were GABAergic; as excitatory HD cells were not targeted, their contributions might have been missed. Second, in contrast to Drosophila, activity was recorded from somata rather than neurites, and somatopetal signal propagation may introduce nonlinearities. Third, the saturation of calcium signals could result in the observed activity plateau. Fourth, the recorded HD cells might receive signals from, rather than constitute, a ring attractor. Fifth, the animals were head-restrained, and AHV was estimated based on tail movement, which might introduce a non-linear velocity-dependent bias. At least in principle, there is also the possibility that the zebrafish HD system implements a more complex ring-attractor network, such as a six-ring structure ^25^. These alternative possibilities will be explored in future research.

The current paper focuses on AHV integration mechanisms and their obligatory consequences for neuronal activity. However, we note that add-on mechanisms might affect activity patterns and blur the distinction between a synapse-modulation network and a shifter network. In fact, by appropriately scaling the overall synaptic strength, we can hand-craft asymmetric AHV tuning in a synapse-modulation network. However, such an ad-hoc mechanism lacks any intrinsic necessity and thus differs fundamentally from the emergent AHV asymmetry characteristic of shifter networks.

Our approach is particularly useful for vertebrate species, such as fish, birds, and mammals, for which identifying the anatomical circuit connectivity is more challenging than in flies. In fact, in the dorsal tegmental nucleus and lateral mammillary nuclei in rats, there are HD cells with no AHV tuning, cells with AHS tuning-only, and cells with skewed AHV tuning ^26^. Our theoretical framework suggests that these two brain regions contain a multi-ring shifter circuit, an indication that a key computational motif for spatial navigation is conserved from fish to rodents. Given that zebrafish and Drosophila diverged at least 550 million years ago, the functional similarity of their shifter networks likely reflects convergent evolution.

## Methods

### Data selection and classification with linear regression

Data from Turner-Evans et al. ^10^ were generously made available by the authors. The data consist of recordings from 14 flies that were head-fixed but could move their legs and abdomen over a free-floating foam ball. Each fly was recorded in a dark or light environment. Within each environment, five one-minute recordings were performed. The calcium activity of GCaMP6f from the protocerebral bridge was used for classification. EPG neurons were divided into 16 ROIs. PEN-L and PEN-R neurons were each divided into eight ROIs. This resulted in a total of 224 EPG, 112 PEN-L, and 112 PEN-R ROIs. See Turner-Evans et al. ^10^ for details. Recordings with visual feedback that rotated two times faster than the flies’ actual rotation were not included in the analysis. In addition to the linear regression described in the main text, the condition (dark and light) and the interaction between the condition and the AHS were used as predictors.

Data from Petrucco et al. ^16^ are publicly available. The dataset consists of volumetric lightsheet imaging in zebrafish larvae expressing GCaMP6s in GABAergic neurons of the anterior hindbrain. Zebrafish were head-restrained but free to move their tails, and recordings were made in darkness or with a visual stimulus in either closed or open-loop conditions. The AHV was estimated by convolving the angles of tail movement bouts with an exponential decay kernel (time constant: three seconds). Four fish that rotated fewer than three times in one direction were excluded, resulting in 27 fish for the analysis. Individual animal IDs correspond to the sorted order of folders in the deposited dataset.

We applied linear regression as described in the main text and classified each ROI or neuron as belonging to the symmetric, CW shifter, or CCW shifter rings. The linear regression coefficient was considered significantly positive (or negative) if the coefficient was greater (or smaller) than zero and the p-value was below 0.05 after applying the Bonferroni correction for multiple comparisons.

### Synapse-modulation network

In ring networks, neurons can be indexed by an angular variable *θ*, − *π ⩽ θ ⩽ π*. Each neuron *θ* has a time-dependent membrane potential *u(θ, t)* and receives input *f(u*(*θ*^′^, *t*)) from other cells at positions *θ*′. The function *f*(*u)* describes the cells’ nonlinear transformation from membrane potential to firing rate. An example of *f (u)* is the piecewise linear *f (u*) = max(*u*, 0), also known as the ReLU (Rectified Linear Unit) function. The synaptic coupling pattern is assumed to be position-invariant, so the coupling strength *W (θ −θ* ′ between two neurons is a function solely of the difference Δ*θ* = *θ −θ*^′^ in the angular index of these two neurons. The neuronal dynamics are

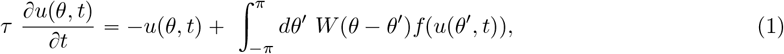

where *τ* is the neuronal membrane time constant. When the synaptic strength is sufficiently strong, spatially modulated, and bidirectional *W (*Δ*θ*) = *W* (−Δ*θ*), i.e., an even function of Δ*θ*, external input or fluctuations can lead to symmetry-breaking and the formation of a persistent and stationary activity bump on the ring. Such a stationary solution is also known as an attractor state.

If the activity bump is to act as a compass, it should move when the animal changes its head direction. Now, if an activity bump moves without changing its shape, it obeys the advection equation ∂ *u*(*θ, t*) /∂ *t* + *ω*(*t*) ∂ *u* (*θ, t*) /∂ *θ* = 0, where *ω*(*t*) is the angular velocity. Zhang ^12^ observed that shifting the ∂ /∂ *θ* operator onto the coupling matrix yields a symmetric and stable traveling solution *u*^*^ (*θ* − *ωt*). In this case, the angular head velocity must modulate the coupling strength as follows:

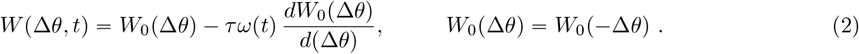

The term *τω*(*t*) *dW*_0_(Δ*θ)* /*d*(Δ*θ*) introduces a time-varying, velocity-dependent asymmetry to the effective couplings, which forces the bump to move.

### Shifter network

A HD shifter network typically consists of three rings of HD neurons, whose activities are denoted by *a*_cw_ (CW shifter ring), *a*_ccw_ (CCW shifter ring), and *a*_sym_ (symmetric ring), respectively. Their dynamics is described by Equations (3) to (8). In some simulations, we omitted the symmetric ring to obtain a two-ring scenario similar to that in Xie et al. ^21^.

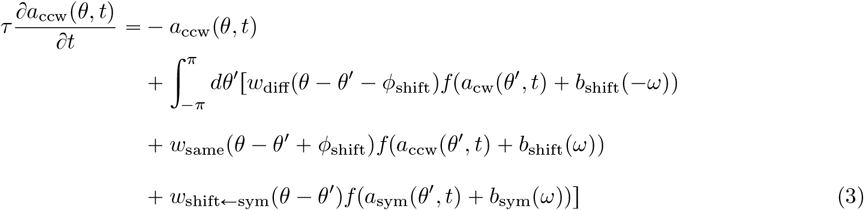

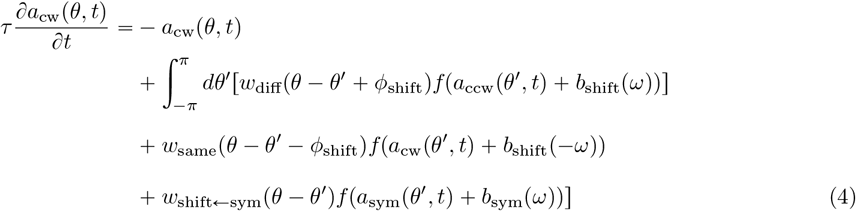

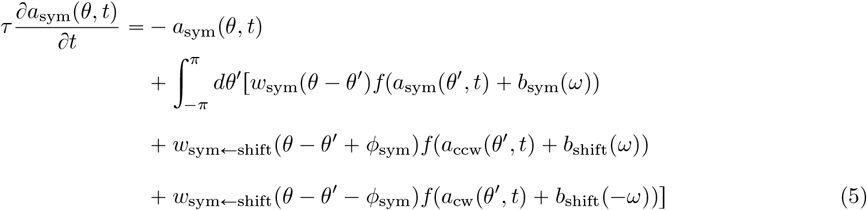

The neuronal activation functions *f* (*a*) of the shifter rings depend on the animal’s AHV via additive bias terms, *b*_shift_(*ω*) and *b*_shift_(− *ω*), for the CCW and CW ring, respectively. Similarly, neurons in the symmetric ring receive a bias *b*_sym_(*ω*) . In all three cases, the time-dependence of *ω* has been omitted for simplicity. As some bias *b* is added to the neuronal activity *a* in the argument of *f*, this is equivalent to a shift of *f* by *a*. In – (5), the variables *a* represent neuronal activities before bias terms are added to result in the membrane potentials *u*, whereas the dynamical variables in Xie et al. ^21^ have been interpreted as synaptic activations.

In the terms *w*(*θ − θ*^′^ ± *α*) the subscripts stand for couplings between the two different shifter rings, within the same shifter rings, within the symmetric ring, and couplings from a shifter ring to the symmetric ring (and in the opposite direction). The parameters *ϕ*_shift_, *ϕ*_sym_ are two constant angles creating skewed connections between the respective rings. We require all couplings *w* (*θ*) to be symmetric and periodic,

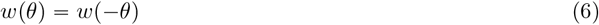

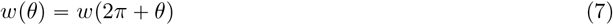

and the odd component of the input *b*_shift_(*ω*) to be proportional to AHV,

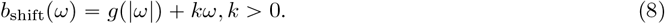

### Rotation-direction preference

To analytically investigate the preference of the shifter rings for CW or CCW rotations, we subtract Eq. (5) from Eq. (4), define *z*(*θ, t*) = *a*_ccw_(*θ, t*) − *a*_cw_(*θ, t*) and obtain:

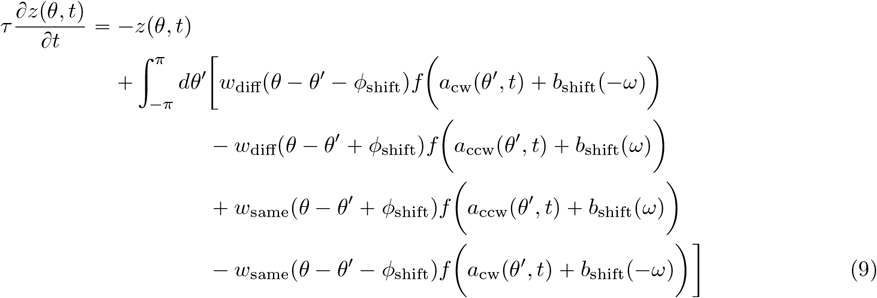

For *w*_diff_ = *w*_same_, Eq. (9) reduces to *τ* ∂*z*(*θ, t*) /∂*t* = −*z*(*θ, t*). In other words, the system relaxes to the solution *z*(*θ, t*) = 0, so that *a*_ccw_(*θ, t*) = *a*_cw_(*θ, t*). In terms of the membrane potentials *u*_ccw_(*θ, t*) = *a*_ccw_(*θ, t*) + *b*_shift_(*ω*) and *u*_cw_(*θ, t*) = *a*_cw_(*θ, t*) + *b*_shift_(− *ω*), Eq.(8) implies

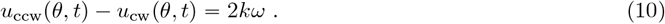

If *w*_diff_ ≠ *w*_same_ we average angular-dependent variables by integrating them from − *π* to *π* and denoting the averaged quantities by < … >. Because the synaptic couplings are 2*π*-periodic, the result of the integration does not depend on the offsets. Eq.(9) becomes:

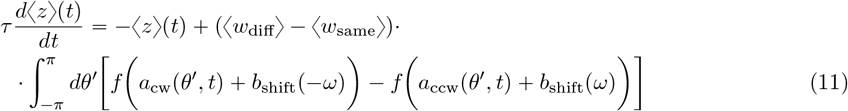

For ⟨ *w*_diff_ ⟩ = ⟨ *w*_same_ ⟩ the stationary solution vanishes, ⟨ *z* ⟩ = 0, i.e., ⟨ *u*_ccw_ ⟩ − ⟨ *u*_cw_ ⟩ = 2*kω*. Otherwise, we simplify the equation and approximate *f*(*x*) by a linear function, *f*(*x*) ≈ *f*(0) + *mx*. Because the activation function is non-decreasing, *m* > 0. Independent of *f* (0) the integration over *θ*^′^ results in

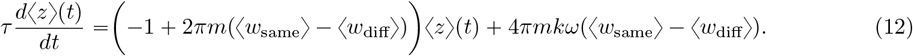

For time-independent *ω*, the stationary solution is given by:

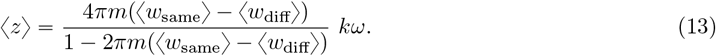

It follows that the difference between the membrane potentials of the CCW and CW rings is

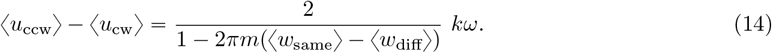

In conclusion, ⟨ *u*_ccw_ ⟩ − ⟨ *u*_cw_ ⟩ is proportional to AHV, although the sign may differ depending on the specific parameters.

Furthermore, *a*_ccw_ (− *θ*, − *ω*) = *a*_cw_(*θ, ω*), *a*_sym_(− *θ*, −*ω*) = *a*_sym_(*θ, ω*) (see Supplementary Information for derivations). Consequently, the average activity difference between the CW and CCW rings is the same as the difference between the CCW and CW activities within one ring. This finding enables the classification of rings based on single-cell activity.

Numerical simulations were employed to test the analytical results, which were performed using the discretized form of the differential equations, where integrals of the form 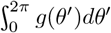 were replaced by 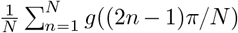 As systems with *N* = 50 or more neurons in each ring showed only small finite-size effects, networks of this size were used throughout the numerical analyses. The *θ* values within each ring were equally distributed in a range from − *π* to *π*. The neuronal time constant *τ* was chosen as 20 ms. A synapse-modulation network (Fig. 1b), a two-ring shifter network (Fig. 1c), and a three-ring shifter network (Fig. 1d) were investigated. Neuronal activation functions *f* (*u*) were either threshold-linear or hyperbolic-tangent functions and synaptic coupling strengths *w* (*θ*) depended on the cells’ HD difference *θ* in the form of cosine or as von Mises functions. The synaptic strength was the same or different for the shifter rings. Consequently, there were four variants for the synapse-modulation network and eight variants for each instance of the shifter-ring network. Grid searches were used for each variant to perform simulations across a large parameter space. To account for the positive modulation of AHS, for the shifter network with two rings, an additional input |*kω*| was added to all cells. For the shifter network with three rings, an additional input |*kω*| . *b*_sym_ was added to the symmetric ring. See Supplementary Information for details.

### Ring-attractor network for the zebrafish HD system

The network is an instance of the three-ring shifter network described by Eqs. (4) to (9), with synaptic couplings given by:

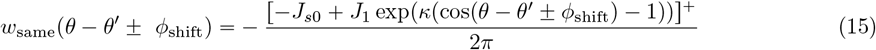

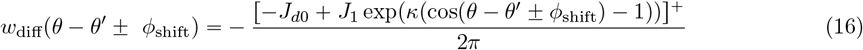

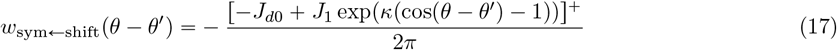

where [ ]^+^ = max(0, *x*) denotes the ReLU threshold-linear function. Angular offsets are taken as *ϕ*_shift_ = − 8 *π* / 9 and *ϕ*_sym_ = 0. To incorporate the abrupt increase and AHS modulation in activity, *b*_shift_ (*ω*) = 1 + 0.1 *ω* + *h*(*ω*), *b*_sym_ (*ω)* = 1 + *h* (*ω)* where *h* (*ω*) = 0.3 . sgn (|*ω*|) + 0.1 . | *ω* |. A grid search was (erformed with *J*_*s*0_, *J*_*d*0_, *J*_1_ ∈ { 0, 10, 20, 30, 40, 50 } and *κ* ∈ { 0.63, 1.10, 1.91, 3.31, 5.75, 10.00 }. All networks that linearly integrated constant AHVs were selected for a test of their performance using real trajectories from the 27 zebrafish. The gain *G* from *k ω* to the moving speed of the activity profile was calculated when we examined whether the networks integrated constant AHVs linearly. When testing the performance using real trajectories, *ω* was the angle measured in every 200-ms time bin of each zebrafish’s trajectory, while the *k* was 1 / *G*. See Supplementary Information for further details.

As proof of principle, a ring attractor network with non-uniformly distributed *θ* was developed:

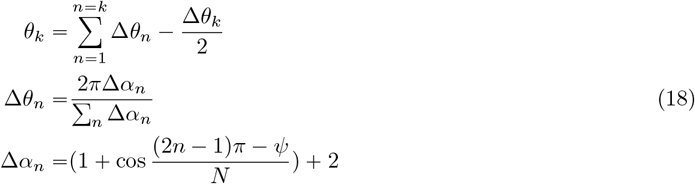

Here, *n* is the index of the neuron and ranges from 1 to *N* = 50. The original synaptic couplings *w* (*θ*_*i*_ − *θ*_*k*_) were scaled by *N* Δ*α*_*n*_ / ∑ _*m*_ Δ*α*_*m*_ to compensate for the non-uniform distribution. *ψ* is the center of the low-density region, which was set to −*π* / 2 for the CCW ring and *π* / 2 for the CW ring.

**Extended Data Fig. 1:**
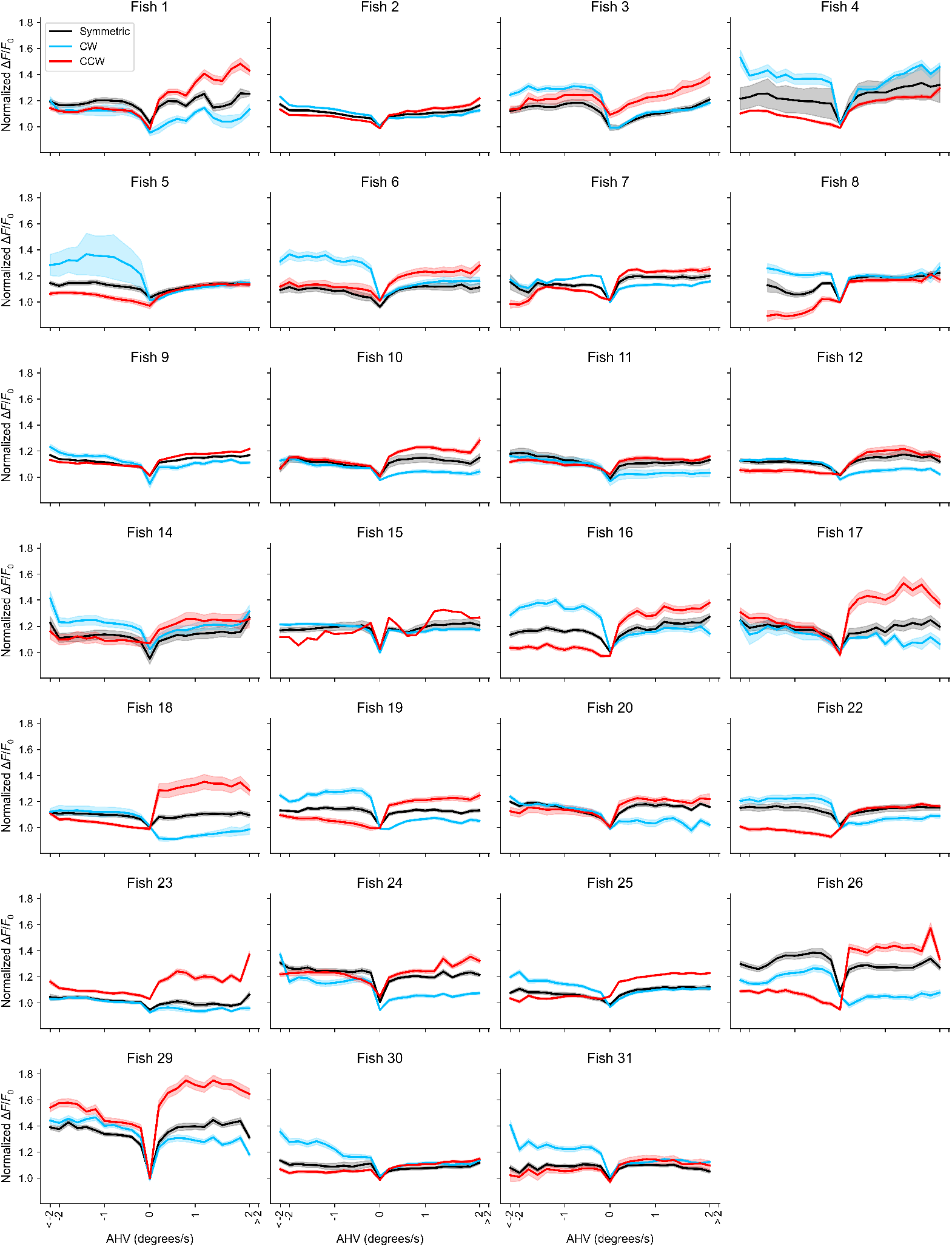
Fish-by-fish data of the neuronal activity as a function of angular head velocity. Shown are the responses of the HD cells assigned to the symmetric ring (in black), CW ring (in blue), and CCW ring (in red), respectively. To isolate the AHV modulation, the variance attributable to HD preference was removed based on linear regression. Averages of these data across fish are depicted in Fig.3f, right panel.

**Extended Data Fig. 2:**
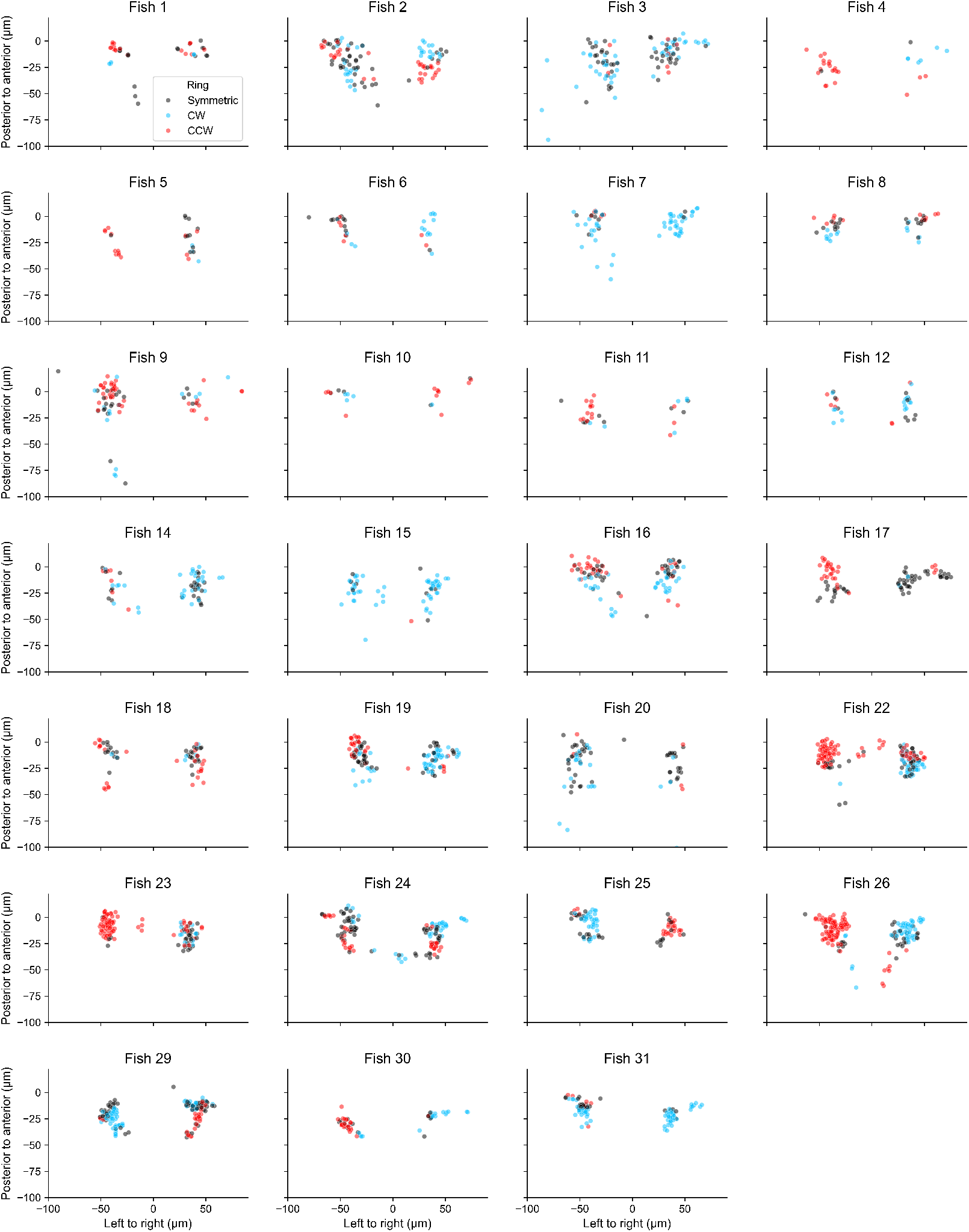
Fish-by-fish data of the anatomical locations of HD cells. As in Fig.3j, which depicts data from all fish, the locations of HD cells assigned to the symmetric ring (in black), CW ring (in blue), and CCW ring (in red), respectively, are shown. HD cells are projected onto the left-to-right vs. posterior-to-anterior plane.

**Extended Data Fig. 3a:**
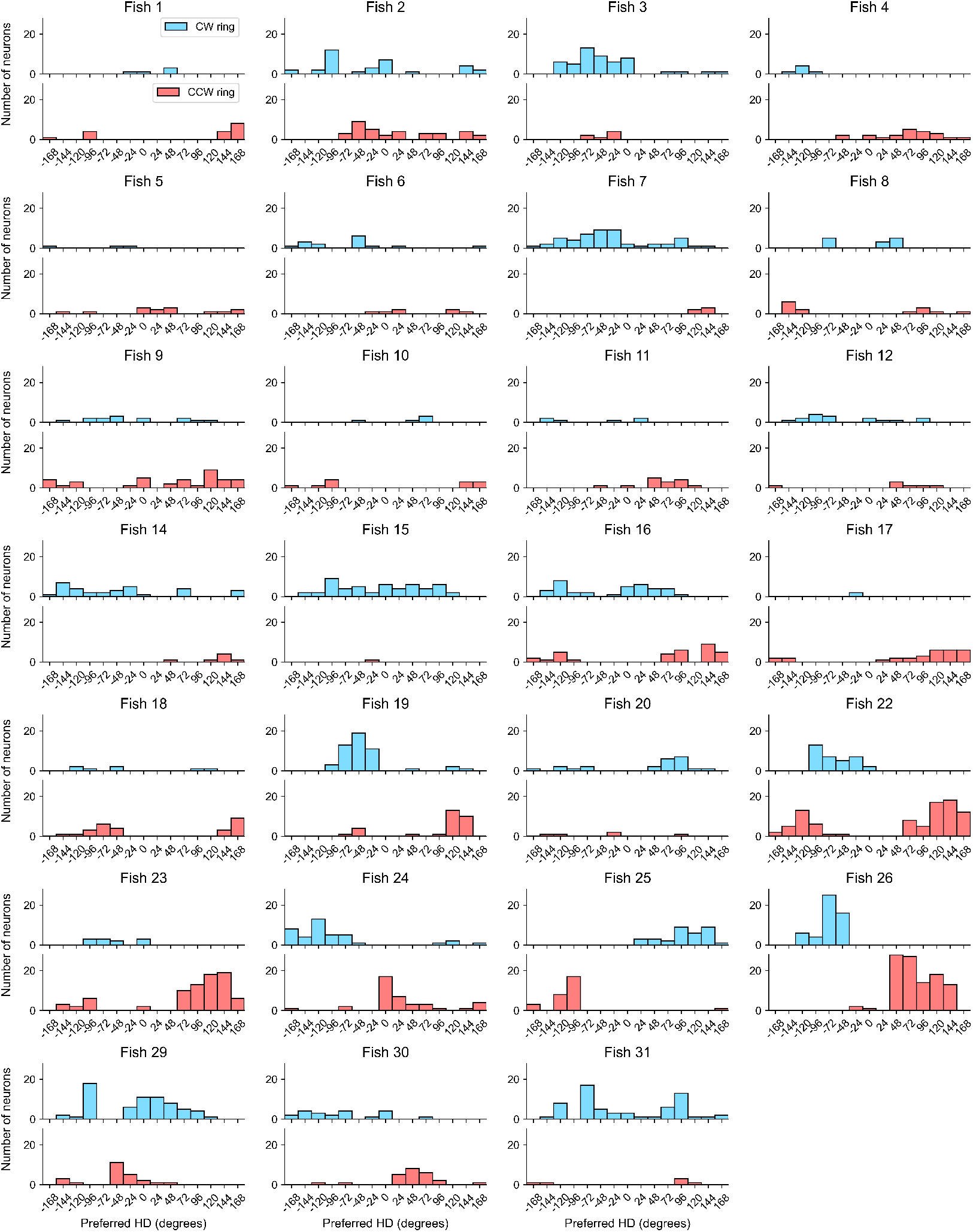
Fish-by-fish data of the distribution of preferred HDs in the shifter rings. As in the population-wide presentation (Fig.3m), data from HD cells assigned to the CW ring (in blue) and to the CCW ring (in red) are shown.

**Extended Data Fig. 3b:**
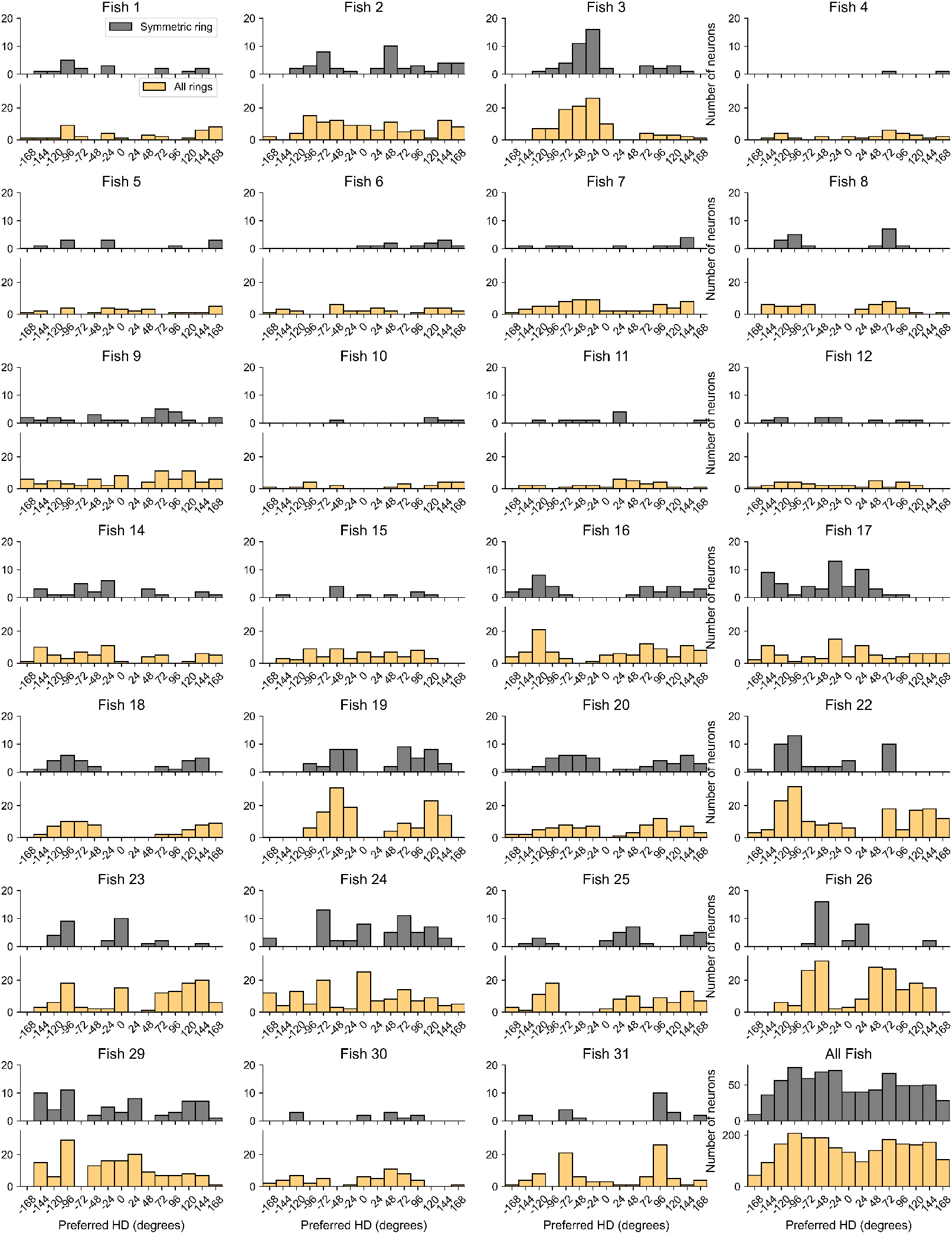
Fish-by-fish data of the distribution of preferred HDs. Complementing Extended Data Fig. 3a, data from HD cells assigned to the symmetric ring (in grey) and from all HD cells (in yellow) are shown.

## Acknowledgements

We are grateful to all members of the Herz and Portugues labs for their input. This research was funded by the German Research Foundation (DFG) under Germany’s Excellence Strategy within the framework of the Munich Cluster for Systems Neurology (EXC 2145 SyNergy, identifier 390857198) and SPP2205 (project 430156228). R.P. was supported by start-up funds from Cornell University.

## Author contributions

The conception and design of the study were done by S.M. with help from all authors. Data analysis, computational modeling, and mathematical analysis were carried out by SM, supported by M.S. and A.V.M.H. The paper was written by S.M., M.S., and A.V.M.H. with contributions from all authors.

