## Supplementary Information for "A multi-ring shifter network computes head direction in zebrafish"

December 2025

### 1 Rotation-direction preference in shifter-ring attractor networks

#### 1.1 Symmetry in AHV $\omega$ and HD $\theta$

In general, the angular velocity $\omega$ is a function $\omega(t)$ of time $t$. The solutions to equations (3) to (8) obey

$$a_{\text{ccw}}(-\theta, -\omega(t)) = a_{\text{cw}}(\theta, \omega(t)), \tag{S1}$$

$$a_{\text{sym}}(-\theta, -\omega(t)) = a_{\text{sym}}(\theta, \omega(t)), \tag{S2}$$

provided that these equations hold true at $t = 0$ as an initial condition. This symmetry is a direct consequence of the synaptic weight couplings obeying $w(\theta - \theta' \pm \alpha) = w(-\theta + \theta' \mp \alpha)$ Implicit in this construction is the assumption that CW and CCW have the same number of neurons, allowing the two shifter rings to balance each other.

Hence, the activity difference between the CW and CCW rings, averaged across $\theta$, is equivalent to the difference between CCW and CW rotations within the same ring.

#### 1.2 Numerical simulations

To numerically simulate the networks, we discretized equations (1) to (8) with $N = 50$ neurons per ring. With this $N$, a reasonably smooth approximation to continuous attractor states was obtained. The neuronal time constant $\tau$ was taken to be 20 ms for all networks. Each neuron is assigned an angular variable $\theta_i$, with the integer index $i$ ranging from 0 to $N - 1$ and the $\theta_i$'s tiling $(-\pi, \pi]$ uniformly. These determined the synaptic couplings, which were either cosine or von Mises functions of the angular variable differences $\theta_i - \theta_j$. Further details on these coupling matrices are given below for the different networks. In most cases, a constant offset was added to the coupling matrix to simulate uniform inhibition.

##### 1.2.1 Synapse-modulation network

A discretized version of equation (1) was implemented:

$$\tau\frac{da_{\theta_i}(t)}{dt} = -a_{\theta_i}(t) + \frac{1}{N}\sum_{j=1}^{N} w(\theta_i - \theta_j, t) f\Big(a_{\theta_j}(t) + b_0\Big) \tag{S3}$$

This network is identical to the one proposed by Zhang[1] since the bias $b_0$ can be integrated in the definition of the neuronal activation function, $f_0(x) = f(x + b_0)$.

To test the influence of different network connectives, we used (a) a cosine synaptic coupling function,

$$\begin{aligned} w(\theta, t) = & J^{\text{offset}} + J^{\text{scale}} \cdot \cos(\theta) \\ & - J^{\text{scale}} \cdot k\omega(t) \cdot \sin(\theta) \ , \end{aligned} \tag{S4}$$

and (b) a von Mises synaptic coupling function,

$$\begin{aligned} w(\theta, t) = & J^{\text{offset}} + J^{\text{scale}} \cdot e^{\kappa(\cos(\theta)-1)} \\ & - J^{\text{scale}} \cdot k\omega(t) \cdot \kappa \sin(\theta) \cdot e^{\kappa(\cos(\theta)-1)} \ . \end{aligned} \tag{S5}$$

Here, $J^{\text{offset}}$ and $J^{\text{scale}}$ are two free parameters, with $J^{\text{offset}} \leq 0$ a constant factor modeling inhibition, and $J^{\text{scale}} \geq 0$ a scaling factor $J^{\text{scale}} \geq 0$. A large (small) concentration parameter $\kappa$ of the von Mises function, results in a narrower (wide) connection range. We performed grid searches over the parameters $J^{\text{offset}}$, $J^{\text{scale}}$ and $\kappa$. Since only the product of $k$ and $\omega$ enter the dynamics, we did not treat these two parametersm individually but rather examined the correlation between the speed of the activity bump and $k\omega$.

We implemented two prominent neuronal activation functions, the threshold linear “ReLU" function,

$$f(x) = \max(0, x) \ , \tag{S6}$$

and the hyperbolic tangent

$$f(x) = \frac{\tanh(x) + 1}{2} \ . \tag{S7}$$

Two synaptic coupling functions and two activation functions resulted in four variants of synapse-modulation networks. Note that a network with a threshold- linear function and zero external input cannot form a ring attractor, because if $a_0(\theta, t)$ is a stable state solution to the network, then for any $c > 0$, $c \cdot a_0(\theta, t)$ is also a solution.

##### 1.2.2 Shifter network

We implemented a discretized version of the shifter network with two rings (Fig. 1c) and three rings (Fig. 1d). For two rings, this amounts to

$$\begin{aligned}\tau \frac{da_{\theta_i}^{\text{ccw}}(t)}{dt} =& - a_{\theta_i}^{\text{ccw}}(t) + \\ & \frac{1}{N}\sum_{j=1}^{N}\Bigg[w_{\text{same}}(\theta_i - \theta_j + \phi_{\text{shift}}, t) f\Bigg( a_{\theta_j}^{\text{ccw}}(t) + b_{\text{shift}}(\omega)\Bigg) + \\ & \quad w_{\text{diff}}(\theta_i - \theta_j - \phi_{\text{shift}}, t) \ f\Bigg( a_{\theta_j}^{\text{cw}}(t) + b_{\text{shift}}(-\omega)\Bigg)\Bigg] \ ,\end{aligned} \tag{S8}$$

$$\begin{aligned}\tau \frac{da_{\theta_i}^{\text{cw}}(t)}{dt} =& - a_{\theta_i}^{\text{cw}}(t) + \\ & \frac{1}{N}\sum_{j=1}^{N}\Bigg[w_{\text{same}}(\theta_i - \theta_j - \phi_{\text{shift}}, t) f\Bigg( a_{\theta_j}^{\text{cw}}(t) + b_{\text{shift}}(-\omega)\Bigg) + \\ & \quad w_{\text{diff}}(\theta_i - \theta_j + \phi_{\text{shift}}, t) f\Bigg( a_{\theta_j}^{\text{ccw}}(t) + b_{\text{shift}}(\omega)\Bigg)\Bigg] \ .\end{aligned} \tag{S9}$$

Here, $N = 50$, $\phi_{\text{shift}} = \pi/9$, $b_{\text{shift}}(\omega) = b_0 + k\omega, b_{\text{shift}}(-\omega) = b_0 - k\omega$, and $b_0$ was chosen as $b = 1$. This network is a special case of the model proposed by[2] though with transformed dynamical variables (see Suppl. Information, Section 4mat. XX).

We analyzed networks with $w_{\text{same}} = w_{\text{diff}}$, and with $w_{\text{same}} \neq w_{\text{diff}}$. For networks with

$w_{\text{same}} \neq w_{\text{diff}}$, synaptic couplings were either chosen as cosine functions,

$$w_{\text{same}}(\theta) = J_{\text{same}}^{\text{offset}} + J^{\text{scale}} \cdot \cos(\theta) \ , \tag{S10}$$

$$w_{\text{diff}}(\theta) = J_{\text{diff}}^{\text{offset}} + J^{\text{scale}} \cdot \cos(\theta) \ , \tag{S11}$$

or von Mises functions,

$$w_{\text{same}}(\theta) = J_{\text{same}}^{\text{offset}} + J^{\text{scale}} \cdot e^{\kappa(\cos(\theta)-1)} \ , \tag{S12}$$

$$w_{\text{diff}}(\theta) = J_{\text{diff}}^{\text{offset}} + J^{\text{scale}} \cdot e^{\kappa(\cos(\theta)-1)} \ . \tag{S13}$$

For networks with $w_{\text{same}} = w_{\text{diff}}$, we set $J_{\text{diff}}^{\text{offset}} = J_{\text{same}}^{\text{offset}}$. Here, $J_{\text{same}}^{\text{offset}}$, $J_{\text{diff}}^{\text{offset}}$, $J^{\text{scale}}$ are three free parameters; $J_{\text{same}}^{\text{offset}}$ is a constant inhibitory background, $J_{\text{same}}^{\text{offset}} \leq 0$; $J_{\text{diff}}^{\text{offset}}$ can be positive or negative, and $J^{\text{scale}}$ scales the excitatory contribution $J^{\text{scale}} \geq 0$. As Xie et al.[2] proved that the scaling factor of $w_{\text{same}}$ and $w_{\text{diff}}$ have to be identical for linear AHV integration, we use the same convention.

For the activation function, we used either a threshold-linear or a hyperbolic tangent function as described by equations (S6) and (S7). In total, the two synaptic coupling functions, two activation functions, and two versions of $w_{\text{same}}$ and $w_{\text{diff}}$ resulted in eight variants of the two-ring shifter network.

Similar considerations apply to the three-ring shifter network, whose dynamical equations are given by

$$
\begin{aligned}
\tau\frac{da^{\text{ccw}}_{\theta_i}(t)}{dt} =& - a^{\text{ccw}}_{\theta_i}(t)+\\
&\frac{1}{N}\sum_{j=1}^{N}\Bigg[w_{\text{diff}}(\theta_i-\theta_j)f\Bigg(a^{\text{cw}}_{\theta_j}(t)+b_{\text{shift}}(-\omega)\Bigg)+\\
&\qquad w_{\text{same}}(\theta_i-\theta_j)f\Bigg(a^{\text{ccw}}_{\theta_j}(t)+b_{\text{shift}}(\omega)\Bigg)+\\
&\qquad w_{\text{shift}\leftarrow\text{sym}}(\theta_i-\theta_j)f\Bigg(a^{\text{sym}}_{\theta_j}(t)+b_{\text{sym}}\Bigg)\Bigg]\ ,
\end{aligned}
\tag{S14}
$$

$$
\begin{aligned}
\tau\frac{da^{\text{ccw}}_{\theta_i}(t)}{dt} =& - a^{\text{ccw}}_{\theta_i}(t)+\\
&\frac{1}{N}\sum_{j=1}^{N}\Bigg[w_{\text{diff}}(\theta_i-\theta_j)f\Bigg(a^{\text{cw}}_{\theta_j}(t)+b_{\text{shift}}(-\omega)\Bigg)+\\
&\qquad w_{\text{same}}(\theta_i-\theta_j)f\Bigg(a^{\text{ccw}}_{\theta_j}(t)+b_{\text{shift}}(\omega)\Bigg)+\\
&\qquad w_{\text{shift}\leftarrow\text{sym}}(\theta_i-\theta_j)f\Bigg(a^{\text{sym}}_{\theta_j}(t)+b_{\text{sym}}\Bigg)\Bigg]\ ,
\end{aligned}
\tag{S15}
$$

$$
\begin{aligned}
\tau\frac{da^{\text{sym}}_{\theta_i}(t)}{dt} =& - a^{\text{sym}}_{\theta_i}(t)+\\
&\frac{1}{N}\sum_{j=1}^{N}\Bigg[w_{\text{sym}}(\theta_i-\theta_j)f\Bigg(a^{\text{sym}}_{\theta_j}(t)+b_{\text{sym}}\Bigg)+\\
&\qquad w_{\text{sym}\leftarrow\text{shift}}(\theta_i-\theta_j+\phi_{\text{sym}})f\Bigg(a^{\text{ccw}}_{\theta_j}(t)+b_{\text{shift}}(\omega)\Bigg)+\\
&\qquad w_{\text{sym}\leftarrow\text{shift}}(\theta_i-\theta_j-\phi_{\text{sym}})f\Bigg(a^{\text{cw}}_{\theta_j}(t)+b_{\text{shift}}(-\omega)\Bigg)\Bigg]\ .
\end{aligned}
\tag{S16}
$$

We set $b_{\text{shift}}(\omega) = b_0 + k\omega, b_{\text{shift}}(-\omega) = b_0 - k\omega, b_0 = 1$, $b_{\text{sym}} = 5$, $\phi_{\text{sym}} = -8\pi/9$. This class of network is reminiscent of the models proposed by Boucheny at al.[3] and Song and Wang[4], as well as the model for the fly HD network[5].

We studied networks with $w_{\text{same}} = w_{\text{diff}}$ and $w_{\text{same}} \neq w_{\text{diff}}$. For networks with $w_{\text{same}} =$

$w_{\text{diff}}$, the synaptic couplings were either cosyne functions,

$$w_{\text{same}}(\theta) = w_{\text{diff}}(\theta) = J^{\text{scale}}_{\text{shift}} \cdot (\cos(\theta) - 1) \ , \tag{S17}$$

$$w_{\text{shift}\leftarrow\text{sym}}(\theta) = J^{\text{scale}}_{\text{shift}\leftarrow\text{sym}} \cdot (1 + \cos(\theta)) \ , \tag{S18}$$

$$w_{\text{sym}\leftarrow\text{shift}}(\theta) = -J^{\text{scale}}_{\text{sym}\leftarrow\text{shift}} \cdot (1 + \cos(\theta)) \ , \tag{S19}$$

or von Mises functions,

$$w_{\text{same}}(\theta) = w_{\text{diff}}(\theta) = J^{\text{scale}}_{\text{shift}} \cdot e^{\kappa(-\cos(\theta)-1)} \ , \tag{S20}$$

$$w_{\text{shift}\leftarrow\text{sym}}(\theta) = J^{\text{scale}}_{\text{shift}\leftarrow\text{sym}} \cdot e^{\kappa(\cos(\theta)-1)} \ , \tag{S21}$$

$$w_{\text{sym}\leftarrow\text{shift}}(\theta) = -J^{\text{scale}}_{\text{sym}\leftarrow\text{shift}} \cdot e^{\kappa(\cos(\theta)-1)} \ . \tag{S22}$$

For networks with $w_{\text{same}} \neq w_{\text{diff}}$, we set $w_{\text{diff}} = w_{\text{shift}\leftarrow\text{sym}} = w_{\text{sym}\leftarrow\text{shift}}$ and $w_{\text{same}} = 0$. The three nonzero scaling parameters, $J^{\text{scale}}_{\text{shift}}$, $J^{\text{scale}}_{\text{shift}\leftarrow\text{sym}}$, $J^{\text{scale}}_{\text{sym}\leftarrow\text{shift}}$, separately control the synaptic strength between the two shifter rings, from the symmetric ring to both shifter rings, and from the two shifter rings to the symmetric ring.

As for the two-ring model, we implemented threshold-linear (S6) or hyperbolic-tangent (S7) functions. Two synaptic-coupling functions, two activation functions and two versions of $w_{\text{same}}$ and $w_{\text{diff}}$ resulted in a total of eight variants of the three-ring shifter network.

##### 1.2.3 Range of parameters

For each type of network, there were four (synapse modulation) or eight (shifter network) model variants, and there were two to four free parameters for each synaptic coupling function. We performed grid searches over these parameters for each model variant. Except for $\kappa$, parameters ranged from 0 to 100. For $\kappa$, based on pilot simulations, we selected the range that resulted in the largest variability in network performance. We used the same parameter values for different networks whenever feasible.

For the synapse-modulation network and for the two-ring shifter network $w_{\text{same}} = w_{\text{diff}}$, we performed grid searches with $J^{\text{offset}} = -4n_0$, $J^{\text{scale}} = 4n_1$, where $n_0, n_1 \in [0, 25] \cap \mathbb{Z}$;

and, if applicable, $\kappa = 10^{-0.6+0.24n_2}, n_2 \in [0,\ 5] \cap \mathbb{Z}$. This choice resulted in 676 networks with cosine couplings and 4056 networks with the von Mises couplings.

For the two-ring shifter network with $w_{\text{same}} \neq w_{\text{diff}}$, we carried out grid searches with $J^{\text{offset}}_{\text{same}} = -10n_0, J^{\text{scale}} = 10n_1$,where $n_0, n_1 \in [0, 10] \cap \mathbb{Z}$; and $J^{\text{offset}}_{\text{diff}} = -100 + 20n_2, n_2 \in [0, 9] \cap \mathbb{Z}$, and, if applicable, $\kappa = 10^{-0.6+0.24n_2}, n_2 \in [0, 5] \cap \mathbb{Z}$. This resulted in 1210 networks with cosine couplings and 7260 networks with von Mises couplings.

For the three-ring shifter model network, we performed grid searches with $J^{\text{scale}}_{\text{sym}\leftarrow\text{shift}} = 10n_0, J^{\text{scale}}_{\text{shift}\leftarrow\text{sym}} = 10n_1$, where $n_0,\ n_1 \in [1, 10] \cap \mathbb{Z}$; $J^{\text{scale}}_{\text{shift}} = 10n_2, n_2 \in [0, 10] \cap \mathbb{Z}$; and, if applicable, $\kappa = 10^{-0.6+0.24n_3}, n_3 \in [0, 5] \cap \mathbb{Z}$. We did not include $J^{\text{scale}}_{\text{sym}\leftarrow\text{shift}} = 0$ and $J^{\text{scale}}_{\text{shift}\leftarrow\text{sym}} = 0$ because the connections between the symmetric ring and the shifter rings should be non-zero to move the activity bump. This setting resulted in 1100 networks with cosine couplings and 6600 networks the von Mises couplings.

If none of the investigated networks linearly integrated the angular head velocity, we carried out additional grid searches with parameters that more densely covered the area that might result in valid networks. This happened once for the three-ring shifter network with $J^{\text{scale}}_{\text{sym}\leftarrow\text{shift}} = J^{\text{scale}}_{\text{shift}\leftarrow\text{sym}}$ and threshold-linear activation. To more closely analyze such networks, we performed an additional grid search with $J^{\text{scale}}_{\text{sym}\leftarrow\text{shift}} = 2n_0$, $J^{\text{scale}}_{\text{shift}\leftarrow\text{sym}} = 2n_1$, where $n_0,\ n_1 \in [1, 10] \cap Z$, and $J^{\text{scale}}_{\text{shift}} = 0.5n_2$, where $n_2 \in [0, 10] \cap Z$, which resulted in 1100 extra networks.

##### 1.2.4 Analysis of HD networks – stable stationary solutions

To test the integration properties of the HD networks for nonzero AHV, we first selected networks that had a stable single bump for zero AHV ($\omega = 0$). To this end, each ring was initialized as $[\cos(\theta) \cdot 10^{-8}]^+ + 10^{-12}$, where $[x]^+ = \max(0, x)$. The small $\cos(\theta)$ term was used to anchor the peak location of the activity profile near $\theta = 0$ and the minute term $10^{-12}$ was included to avoid division by zero. The Python function "scipy integrate.solve_ivp" in "scipy 1.9.1" was used to solve the differential equations, employing an explicit Runge-Kutta method of order 5(4). We ran simulations for networks with different parameters separately. Each simulation progressed iteratively, with numerical integration performed over sequential examination intervals: $0-300$ ms, $300-400$ ms, $400-500$ ms, and so on. After each interval, the dynamical system was examined for convergence to a stable state. The process continued until convergence was confirmed or the simulation time reached 1000 ms. The initial interval ($0-300$ ms) was longer than the following ones to allow the network to evolve to an approximately stable state before being examined for the first time. The value of $a_{\theta_i}(t)$ was measured every 50 ms. The first and the last recorded time points for each examination interval were denoted by $t_0$ and $t_{\text{end}}$, respectively.

For each examination interval, we first determined the precision of integration. The numerical integration kept the local error estimates less than $10^{-6} \times |a_{\theta_i}^{\text{ring}}(t)| + a_{\text{tol}}$ with $a_{\text{tol}} = \min_{\text{ring}}(\max_{\theta_i}(a_{\theta_i}^{\text{ring}}(t_0)) - \min_{\theta_i}(a_{\theta_i}^{\text{ring}}(t_0))) \times 10^{-6}$. We terminated the integration if the maximum of any $a_{\theta_i}^{\text{ring}}(t)$ exceeded $10^9$.

At the end of an examination interval, we checked whether the network formed one stable activity bump for each ring. First, we assessed whether a bump had formed at all. We defined $\delta a_{\theta_i}^{\text{ring}}(t) = a_{\theta_i}^{\text{ring}}(t) - \min_{\theta_i}(a_{\theta_i}^{\text{ring}}(t))$ and checked whether

$$\overline{\delta a_{\theta_i}}^{\text{ring}}(t_{\text{end}}) < 10^{-3} \tag{S23}$$

and

$$\left|\overline{\delta a_{\theta_i}}^{\text{ring}}(t_{\text{end}}) - \delta a_{\theta_i}^{\text{ring}})\right| / \left|\overline{\delta a_{\theta_i}}^{\text{ring}}(t_{\text{end}})\right| < 0.01 \tag{S24}$$

where

$$\overline{\delta a_{\theta_i}}^{\text{ring}}(t_{\text{end}}) = \frac{1}{N}\sum_{i=1}^{N} \delta a_{\theta_i}^{\text{ring}}(t_{\text{end}}) \tag{S25}$$

If these conditions were met for all rings and $\theta_i$, we concluded that no bump had emerged and terminated the simulation.

Next, we checked whether the network formed multiple peaks for each ring. A peak in a ring was defined as a local maximum at $\theta_j$ and had to meet one of the following conditions:

$$\delta a^{\text{ring}}_{\theta_i}(t_{\text{end}}) > 0.3 \max_{\theta_i}(\delta a^{\text{ring}}_{\theta_i}(t_{\text{end}})) \tag{S26}$$

or

$$\delta a^{\text{ring}}_{\theta_i}(t_{\text{end}}) - \delta a^{\text{ring}}_{\theta_i,0}(t_{\text{end}}) > 0.1 \max_{\theta_i}(\delta a^{\text{ring}}_{\theta_i}(t_{\text{end}})) \tag{S27}$$

Here, $\delta a^{\text{ring}}_{\theta_i,0}(t_{\text{end}})$ denotes the activity at the lowest contour of neuron $\theta_i$ and was calculated in three steps: (a) A horizontal line from $\delta a^{\text{ring}}_{\theta_i}(t_{\text{end}})$ to the left and right was extended until the line either reached $\pi$, $-\pi$, or intersected the activity of another neuron $\theta_j$, where $\delta a^{\text{ring}}_{\theta_j}(t_{\text{end}}) > \delta a^{\text{ring}}_{\theta_i}(t_{\text{end}})$. (b) On each side, the minimal activity below the line was identified. These points were bases. (c) The higher of the two bases was marked as the lowest contour. If multiple peaks were identified, the simulation was terminated.

Third, we asked whether the represented head direction, calculated as vector average, was stable. We defined vectors as $\vec{\delta a}^{\text{ring}}_{\theta_i}(t)$, whose magnitude was $\delta a^{\text{ring}}_{\theta_i}(t)$ and whose direction was $\theta_i$. We checked whether $\overline{\vec{\delta a}_{\theta_i}}^{\text{ring}}(t)$, the vector average of $\vec{\delta a}^{\text{ring}}_{\theta_i}(t)$, averaged across $\theta_i$, changed between $t_{\text{end}}$ and $t_{\text{end}-1}$:

$$\left| \angle\overline{\vec{\delta a}_{\theta_i}}^{\text{ring}}(t_{\text{end}}) - \angle\overline{\vec{\delta a}_{\theta_i}}^{\text{ring}}(t_{\text{end}-1}) \right| > \left( 5 - \frac{4 \cdot (t_{\text{end}} - 200\,\text{ms})}{1000\,\text{ms} - 200\,\text{ms}} \right) \cdot \frac{2\pi}{N} \tag{S28}$$

If this was true for at least one ring, we regarded the activity bump as unstable and terminated the simulation. The right side of the equation was designed to make the condition increasingly strict as the time approached 1000 ms.

Fourth, we checked whether the population profile kept the same shape between $t_{\text{end}}$ and $t_{\text{end}-1}$:

$$|a^{\text{ring}}_{\theta_i}(t_{\text{end}-1}) - a^{\text{ring}}_{\theta_i}(t_{\text{end}})| < a_{\text{tol}} \tag{S29}$$

Here,

$$a_{\text{tol}} = 0.01 \min(\delta a^{\text{ring}}_{\theta_{\max}}, \overline{|a_{\theta_i}}^{\text{ring}}|) \ , \tag{S30}$$

$$\delta a^{\text{ring}}_{\theta_{\max}} = \max_i(\delta a^{\text{ring}}_{\theta_i}(t_{\text{end}})) \ , \tag{S31}$$

and

$$\overline{|a_{\theta_i}}^{\text{ring}}| = \frac{1}{N}\sum_{i=1}^{N} |a^{\text{ring}}_{\theta_i}(t_{\text{end}})| \ . \tag{S32}$$

If this was true for every ring and $\theta_i$, we regarded the shape of the activity profile as stable.

Fifth, we checked whether the represented direction was strictly stable:

$$\left| \angle \overline{\vec{\delta a}_{\theta_i}}^{\text{ring}}(t_{\text{end}-1}) - \angle \overline{\vec{\delta a}_{\theta_i}}^{\text{ring}}(t_j) \right| < \frac{2\pi}{N}, \ \forall t_j - t_{\text{end}} > -200 \text{ ms} \tag{S33}$$

If it was true for all rings and $\theta_i$, we considered the represented direction stable.

If the represented direction and the population profile were stable, we concluded that the network had reached its stable state and checked whether the activity had formed a bump:

$$\frac{|\overline{\delta a_{\theta_i}}^{\text{ring}}(t_{\text{end}}) - \delta a^{\text{ring}}_{\theta_i}(t_{\text{end}})|}{|\overline{\delta a_{\theta_i}}^{\text{ring}}(t_{\text{end}})|} > 0.1 \tag{S34}$$

If this was true for at least one $\theta_i$ of a ring, we termed this network a "stationary-valid network". Otherwise, we regarded the network as invalid. Only stationary-valid networks were selected for the simulation with $\omega \neq 0$.

##### 1.2.5 Analysis of HD networks – moving activity bumps

Next, we examined whether the angular velocity of the represented head direction correlated linearly with $k\omega$. We performed simulations with $k\omega \in \{-1.0, -0.6, -0.3, -0.1, 0.1,$

0.3, 0.6, 1.0}. For each network with a specific parameter combination, we ran eight simulations for every $k\omega$ value. For the initial state, the network's final state in the stationary state simulation was chosen. The procedure was the same as that in section 1.2.4 if not mentioned otherwise. The examination interval was $0-200$ ms, $200-300$ ms, $300-400$ ms, and so on, until it reached 1000 ms. We set the first examination interval shorter than in the simulations of the stationary state because pilot simulations suggested that 200 ms are enough to reach a stable state in most cases. The network state was sampled every 20 ms, resulting in a maximum calculable velocity of 25 turns (or 9000 degrees) per second. We denote $t_j$ as any recorded time point of the current examination interval.

At the end of each examination interval, we checked whether the population profile was flat or had multiple peaks. Next, we computed the angular velocity of the represented direction, $\omega_{\text{neural}}$, for each ring, based on the angle of the vector average, $\angle\overrightarrow{\delta a_{\theta_i}}^{\text{ring}}(t_j)$. First, we computed the draft speed as the difference in the represented direction between two consecutively recorded time points:

$$\omega_{\text{draft}}^{\text{ring}}(t_j) = \left[\angle\overrightarrow{\delta a_{\theta_i}}^{\text{ring}}(t_j) - \angle\overrightarrow{\delta a_{\theta_i}}^{\text{ring}}(t_{j-1})\right] \cdot \frac{N}{2\pi} \tag{S35}$$

whose unit is neurons per 20 ms or turns per second (as $N = 50$). Next, we calculated the maximum direction difference (unit: one neuron) in the current examination interval:

$$d_{\text{max}}^{\text{ring}} = \max_j \left|\angle\overrightarrow{\delta a_{\theta_i}}^{\text{ring}}(t_{\text{end}}) - \angle\overrightarrow{\delta a_{\theta_i}}^{\text{ring}}(t_j)\right| \cdot \frac{N}{2\pi} \tag{S36}$$

If for either ring, $d_{\text{max}}^{\text{ring}} < 1$ and the signs of $\omega_{\text{draft}}^{\text{ring}}(t_j)$ were not the same across $t_j$ and/or $d_{\text{max}}^{\text{ring}} < 0.01$, then we considered the activity bump as stationary and terminated the simulation. Otherwise, we regarded the rotation direction as the majority of the signs of $\omega_{\text{draft}}^{\text{ring}}(t_j)$. We extend the range of $\angle\overrightarrow{\delta a_{\theta_i}}^{\text{ring}}(t_j)$ from $[-\pi, \pi)$ to $[-\pi, \infty)$ by adding $2\pi$ to $\angle\overrightarrow{\delta a_{\theta_i}}^{\text{ring}}(t_j)$ whenever the sign of $\omega_{\text{draft}}^{\text{ring}}(t_j)$ was not the same as the rotation direction. The resultant variable was $\theta^{\text{ring}}(t_j)$. Next, we calculated the true angular velocity $\omega^{\text{ring}}(t_j) = N(\theta^{\text{ring}}(t_j) - \theta^{\text{ring}}(t_{j-1}))/2\pi$. We additionally assessed whether $\omega^{\text{ring}}(t_j)$ exceeded the maximum calculable speed and found that none of $\omega^{\text{ring}}(t_j)$ was larger than eight turns per second (or 2880°/s).

Finally, we examined whether the angular velocity, $\omega^{\text{ring}}(t_j)$, was stable for each ring. First, we checked whether:

$$|\max(\omega^{\text{ring}}(t_j)) - \min(\omega^{\text{ring}}(t_j))| < 0.2 \text{ turns per second} \tag{S37}$$

Second, we aligned the population profiles $a^{\text{ring}}_{\theta_i}(t_j)$ on the $\theta$ axis by shifting them by $R[n_j\overline{\omega(t_j)}^{\text{ring}}]$ neurons, where $n$ is the number of recorded time points from the start of the examination interval to $t_j$, $\overline{\omega(t_j)}^{\text{ring}}$ is the average angular velocity across all time points, and $R[x]$ rounds $x$ to the nearest integer. The aligned population profiles were $a^{\text{ring}}_{\theta_i,\text{align}}(t_j)$. We checked whether the population profile maintained the same shape:

$$\max_j(a^{\text{ring}}_{\theta_i,\text{align}}(t_j)) - \min_j\,(a^{\text{ring}}_{\theta_i,\text{align}}(t_j)) < \max_i \left| a^{\text{ring}}_{\theta_i,\text{align}}(t_{\text{end}}) - a^{\text{ring}}_{\theta_{i-1},\text{align}}(t_{\text{end}}) \right| \tag{S38}$$

If the above two conditions were met for all rings and all $\theta_i$, we regarded the angular velocity as stable and terminated the simulation. The represented angular velocity was calculated as $\omega_{\text{neural}}(\omega) = \frac{1}{Mn_{\text{ring}}}\sum_{j=1}^{M}\sum_{\text{ring}}\omega^{\text{ring}}(t_j)$, where $n_{\text{ring}}$ is the number of rings, and $\omega$ is the parameter of the current simulation. If the time reached 1000 ms and the above conditions were not met, the simulation was terminated and the network was considered unstable.

The networks with stable angular velocity were further examined for the linearity between $k\omega$ and $\omega_{\text{neural}}(\omega)$. We defined the linear range of a network as $[-k\omega_M, k\omega_M]$. Here, $k\omega_M$ represents the maximum $k\omega$ such that, for any $k\omega_m \leq k\omega_M$, the absolute correlation between the sequence $(-k\omega_m, -k\omega_{m-1}, ..., k\omega_{m-1}, k\omega_m)$ and the corresponding angular velocity of the network ($\omega_{\text{neural}}$) is greater than 0.99. Here, $m, M \in [2, 3, 4] \cup \mathbb{Z}$, $k\omega_0 = 0$, $k\omega_1 = 0.1$, $k\omega_2 = 0.3$, $k\omega_3 = 0.6$, $k\omega_4 = 1.0$. If $k\omega_M = 1$, we regarded these networks to be linearly integrating AHV.

##### 1.2.6 Validity of the mathematical approximations

Here, we examined whether the mathematical predictions apply also to networks with a finite number of neurons and whether the approximations used in the derivation hold. All the examinations were performed for the $a_{\theta_i}(t_{\mathrm{end},\omega})$, where each $t_{\mathrm{end},\omega}$ is the final time point of a simulation with a particular $k\omega$. In this section, we use $a_{\theta_i}(\omega)$ to denote it. For the synapse-modulation networks that are valid in the stationary state, we examined whether the activity profile maintained the same shape regardless of $k\omega$. First, we calculated the unit difference $\Delta a$ as the maximal activity difference between two neighboring neurons in the steady state:

$$\Delta a = \max_{\theta_i} \left| a_{\theta_i}(\omega = 0) - a_{\theta_{i-1}}(\omega = 0) \right| \tag{S39}$$

$\Delta a$ serves as a measure for the imperfect alignment resulting from a finite number of neurons. Next, we align the peak location of the activity bump in the moving-state simulation to zero, obtaining $a_{\theta_i,\mathrm{align}}(\omega)$. In the stationary state, the peak location was already at zero. Then, we calculated the absolute difference between the moving-state activity and the steady-state activity, $d_{\theta_i}(\omega) = |a_{\theta_i,\mathrm{align}}(\omega) - a_{\theta_i}(0)|$, and divided thsis quantity by $\Delta a$ to obtain a normalized deviation: $d_{\theta_i,\mathrm{norm}}(\omega) = d_{\theta_i}(\omega)/\Delta a$. If the normalized deviation exceeded unity, we considered the shape to be mismatched.

The alignment was done in three steps. First, we shifted the activity by $R[\angle\overrightarrow{\delta a_{\theta_i}}(\omega)\frac{N}{2\pi}]$ neurons and got $a_{\theta_i,1}(\omega)$, where $R[x]$ rounds $x$ to the nearest integer. Next, we shifted $a_{\theta_i,1}(\omega)$ toward the positive or negative direction by one neuron, and obtained $a_{\theta_i,2}(\omega)$ and $a_{\theta_i,3}(\omega)$, respectively. Finally, we calculated the relative deviation of $a_{\theta_i,1}(\omega)$, $a_{\theta_i,2}(\omega)$, $a_{\theta_i,3}(\omega)$, and selected the one with the smallest mean relative deviation $\overline{d_{\theta_i,\mathrm{norm}}}(\omega)$ as $a_{\theta_i,\mathrm{align}}(\omega)$.

We additionally assessed whether the peak firing rate $f(a_{\theta_{\max}}(\omega) + b_0)$ (abbreviated as $f_{\theta_{\max}}(\omega)$) was the same regardless of $k\omega$. Here, $\theta_{\max}$ is the $\theta_i$ that produces the maximum $f_{\theta_i}(\omega)$. First, to roughly estimate the error caused by the finite number of neurons, we calculated

$$\Delta f = \frac{1}{2}\max(|f_{\theta_{\max}}(0) - f_{\theta_{\max-1}}(0)|, |f_{\theta_{\max}}(0) - f_{\theta_{\max+1}}(0)|) \tag{S40}$$

We further calculated the normalized maximum firing rate difference,

$$d_{f,\mathrm{norm}}(\omega) = |f_{\theta_{\max}}(\omega) - f_{\theta_{\max}}(0)|/f_{\theta_{\max}}(0)\ , \tag{S41}$$

and examined whether $d_{f,\mathrm{norm}}(\omega) < \frac{\Delta f}{f_{\theta_{\max}}(0)} + 0.01$. We added 0.01 because when there were more than three approximately equal valued maxima, $\Delta f$ was almost zero. If the condition was not met, we considered that the peak firing rate did not match.

For the shifter networks, we examined whether the activity difference between the shifter rings was proportional to $k\omega$. We calculated correlations between $k\omega$ and the difference between the peak firing rate of the CCW and CW rings,

$$f(a^{\mathrm{ccw}}_{\theta_{\max}}(\omega) + b_{\mathrm{shift}}(\omega)) - f(a^{\mathrm{cw}}_{\theta_{\max}}(\omega) + b_{\mathrm{shift}}(-\omega))\ , \tag{S42}$$

and the difference between the mean firing rate of the CCW and CW rings, i.e.,

$$\frac{1}{N}\sum_i [f(a^{\mathrm{ccw}}_{\theta_i}(\omega) + b_{\mathrm{shift}}(\omega)) - f(a^{\mathrm{cw}}_{\theta_i}(\omega) + b_{\mathrm{shift}}(-\omega))]\ . \tag{S43}$$

To calculate these correlations, we included all networks with a stable angular velocity but restricted the range of $k\omega$ to be within its linear range $[-k\omega_i, k\omega_i]$.

We also examined whether the activity of the CW and CCW rings was mirror-symmetric to each other for all networks with a stable angular velocity. We first flipped the activity of the CCW ring. Next, we aligned the activity of the CW and CCW rings by matching their vector average as precisely as possible. Finally, we calculated $d_{\theta_i,\mathrm{norm}}(\omega)$ and $\Delta a$. The procedure was the same as what we did for the synapse-modulation network, with the activity of the CW ring playing the role of the activity of the synapse-modulation network when $\omega = 0$. We also examined the symmetry of the symmetric ring using the same procedure for the shifter network with three rings. When $w_{\mathrm{same}} = w_{\mathrm{diff}}$, we additionally examined whether $a^{\mathrm{ccw}}_{\theta_i}(\omega) = a^{\mathrm{cw}}_{\theta_i}(\omega)$ by verifying that $|a^{\mathrm{ccw}}_{\theta_i}(\omega) - a^{\mathrm{cw}}_{\theta_i}(\omega)| < 10^{-8}$ for all $\theta_i$, $\omega$, and networks that were valid in the stationary state.

#### 1.3 Simulation results: synapse-modulation network

Zhang (1996) theoretically established that the synapse-modulation network keeps the same shape of the activity profile regardless of AHV. We tested this prediction using numerical simulations in a wider parameter space. For all simulated networks, and tested values of $\theta_i$ and $\omega$, the activity matched, $d_{\theta_i,\mathrm{norm}}(\omega) < 1$, $M = 0.103$, $SD = 0.115$, Max $= 0.591$. The peak firing rate also remained the same: all $d_{f,\mathrm{norm}}(\omega) < \Delta f / f_{\theta_{\max}}(0) + 0.01$, $M = 0.005$, $SD = 0.007$, Max $= 0.052$. See supplementary table 1 for more details. To conclude, 3628 out of 9464 simulated synapse-modulation networks exhibited a valid activity bump in the stationary state; all of these networks integrated AHV linearly.

#### 1.4 Simulation results: shifter networks

29541 out of 84484 simulated shifter networks exhibited a valid activity bump when in the stationary state, and 11074 of these shifter networks can integrate AHV linearly.

For a shifter network described by equations (3) to (8), the activity difference between the two shifter rings increases with AHV. In the simulated systems, the mean and peak firing rate differences between the CCW and CW rings increased almost linearly with $\omega$. Mean firing rate: $r : M = 0.993$, $SD = 2.14 \times 10^{-2}$, Min $= 0.566$; peak firing rate: $r : M = 0.999$, $SD = 8.41 \times 10^{-3}$, Min $= 0.323$. See supplementary table 2 for more details. Additionally, if the synaptic coupling function of the two shifter rings is the same ($w_{\mathrm{diff}} = w_{\mathrm{same}}$), then the activity variable $a$ of the CCW and CW rings is exactly the same in the steady state, up to machine precision, $|a^{\mathrm{ccw}}_{\theta_i}(\omega) - a^{\mathrm{cw}}_{\theta_i}(\omega)| < 1.40 \times 10^{-13}$.

The activity of the two shifter rings did not deviate from symmetry in $\theta$ and $\omega$. $d_{\theta_i,\mathrm{norm}}(\omega)$: $M = 2.19 \times 10^{-5}$, $SD = 3.28 \times 10^{-4}$, Max $= 5.20 \times 10^{-2}$. The same holds true for the intrinsic symmetry (regarding $\theta$ and $\omega$) of the symmetric ring. $d_{\theta_i,\mathrm{norm}}(\omega)$: $M = 1.60 \times 10^{-6}$, $SD = 1.60 \times 10^{-4}$, Max $= 4.34 \times 10^{-2}$.

### 2 Positive AHS Modulation

Here, we describe one way the synapse-modulation network can be positively modulated by AHS.

#### 2.1 Methods

The procedure was the same as stated in section 1.2, except for differences in parameters. For the synapse-modulation network, we implemented Eq. (S3) with $f(x) = \max(0, x)$ and $w(\theta) = [J^{\mathrm{offset}}_{\mathrm{same}} + J^{\mathrm{scale}} \cdot \cos(\theta) - J^{\mathrm{scale}} \cdot k\omega \cdot \sin(\theta)](1 - \frac{|k\omega|}{2})$. In other words, we assumed that, in addition to the AHV modulation, increasing the AHS decreases synaptic strength. We performed a grid search with $J^{\mathrm{offset}}_{\mathrm{same}} \in [-100, -98, -96, ..., 0]$ and $J^{\mathrm{scale}} \in [0, 2, 4, ..., 100]$.

For the two-ring shifter network, Eq. (S8) and (S9) with $w_{\mathrm{diff}} \neq w_{\mathrm{same}}$, cosine couplings (S10) and (S11), and $f(x) = \max(0, x)$ were used. Additionally, $b_{\mathrm{shift}}(\omega)$ increased with increasing AHS, $b_{\mathrm{shift}}(\omega) = b_0 + k\omega + |k\omega|$, $b_{\mathrm{shift}}(-\omega) = b_0 - k\omega + |k\omega|$. A grid search with $J^{\mathrm{offset}}_{\mathrm{same}} \in [-100, -90, -80, ..., 0]$, $J^{\mathrm{scale}} \in [0, 10, 20, ..., 100]$ and $J^{\mathrm{offset}}_{\mathrm{diff}} \in [-100, -80, -60, ..., 80]$ was performed.

For the three-ring shifter network, we implemented equations (S14) to (S16) with $w_{\mathrm{same}} \neq w_{\mathrm{diff}}$, cosine couplings eqrefeq:samecos and (S11), and $f(x) = \max(0, x)$. Additionally, we made $b_{\mathrm{sym}}$ increases as the AHS increases: $b_{\mathrm{sym}} = 5 \cdot (1 + |k\omega|)$. We performed a grid search with $J^{\mathrm{scale}}_{\mathrm{sym} \leftarrow \mathrm{shift}} \in [2, 4, 6, ..., 20]$, $J^{\mathrm{scale}}_{\mathrm{shift} \leftarrow \mathrm{sym}} \in [2, 4, 6, ..., 20]$ and $J^{\mathrm{scale}}_{\mathrm{shift}} \in [0, 0.5, 1, ..., 5]$.

We ran the numerical simulation as described in section 1.2.4 and 1.2.5. To examine the effect of AHS on neuronal activity, we applied two linear regression modes, each with a free intercept, using the positive half of $k\omega$ as the predictor and the mean and peak firing rates as the target variables. For the shifter network, we predicted the average ring activity. The magnitude of increase was assessed using the slope of the linear regression, and the ratio of increase was assessed by dividing the slope by the intercept. We also examined whether the theoretical predictions hold, as described in section 1.2.6.

#### 2.2 Results

For the synapse-modulation network, 366 among 676 networks integrate angular head velocity in a linear manner. The firing rate of 317 networks increases as the AHS increases. The deviation of the neuronal activity from symmetry is negligible. $d_{\theta_i,\text{norm}}(\omega) : M = 2.07 \times 10^{-4}, SD = 9.45 \times 10^{-4}, \text{Max} = 2.23 \times 10^{-2}$.

For the two-ring shifter network, 129 out of 1210 networks integrate AHV linearly, and all of these networks exhibit an increased firing rate with increasing AHS. The correlations between the mean and peak firing rate difference of shifter rings and $k\omega$ are almost linear, $M \geq 0.999, SD \leq 4.48 \times 10^{-4}, \text{Min} = 0.996$. The activities of the shifter rings are symmetric with each other regarding $\theta$ and $\omega$, $M = 5.98 \times 10^{-6}, SD = 6.48 \times 10^{-5}, \text{Max} = 4.11 \times 10^{-3}$.

For the three-ring shifter network, 139 out of 1100 networks integrate AHV linearly , and 133 networks show an increased firing rate. The correlations between the mean and peak firing rate difference of shifter rings and $k\omega$ are almost linear, $M \geq 0.996, SD \leq 2.43 \times 10^{-2}, \text{Min} = 0.680$. The activities of the shifter rings are symmetric to each other regarding $\theta$ and $\omega$, $d_{\theta_i,\text{norm}}(\omega)$: $M = 1.18 \times 10^{-6}, SD = 2.02 \times 10^{-5}, \text{Max} = 1.17 \times 10^{-3}$, and the activity of the symmetric ring is symmetric to itself in $\theta$ and $\omega$, $d_{\theta_i,\text{norm}}(\omega)$: $M = 5.45 \times 10^{-7}, SD = 1.30 \times 10^{-5}, \text{Max} = 8.94 \times 10^{-4}$.

### 3 Simulation results: Zebrafish HD system

For networks with uniformly distributed $\theta$, 246 out of the 1296 examined networks integrate AHV linearly. This number decreased to 54 for networks with non-uniformly distributed $\theta$.

The correlations between $k\omega$ and the firing rate difference between shifter rings were close to unity, regardless of whether the HDs were uniformly distributed or not. Mean firing rate: $M = 0.999$, $SD = 9.82 \times 10^{-3}$, $\text{Min} = 0.828$; Peak firing rate: $M = 0.999$, $SD = 2.08 \times 10^{-3}$, $\text{Min} = 0.980$. The activity of the two shifter rings was symmetric in $\theta$ and $\omega$, again regardless of $\theta$ uniformity. $d_{\theta_i,\text{norm}}(\theta)$: $M = 9.00 \times 10^{-7}$, $SD = 2.00 \times 10^{-5}$, Max

$= 1.12 \times 10^{-3}$. The activity of the symmetric ring was symmetric to itself in $\theta$ and $\omega$. $d_{\theta_i,\mathrm{norm}}(\theta)$: $M = 4.15 \times 10^{-7}$, $SD = 1.34 \times 10^{-5}$, $\mathrm{Max} = 9.50 \times 10^{-4}$. See supplementary table 3 for details.

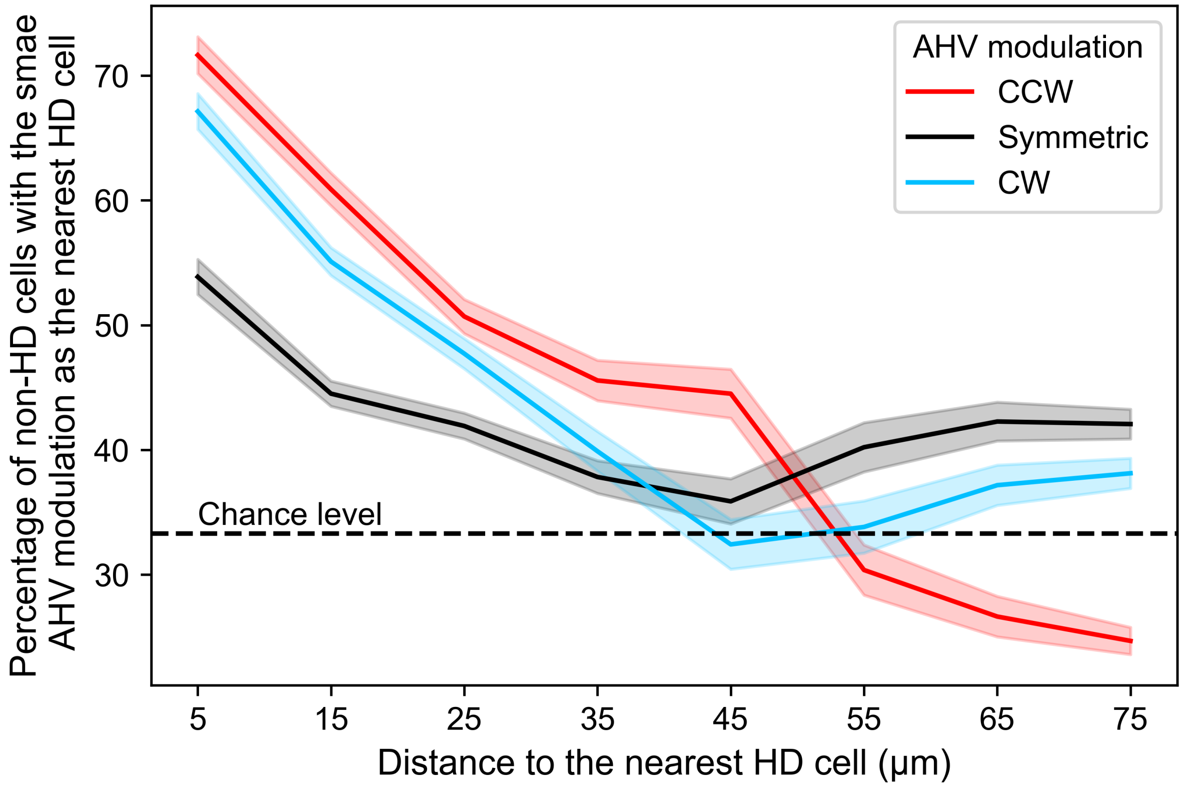


**Fig. S1** | Percentage of non-HD cells that share the same AHV modulation as the nearest HD cells. The percentage decreases as the distance to the nearest HD cells increases.

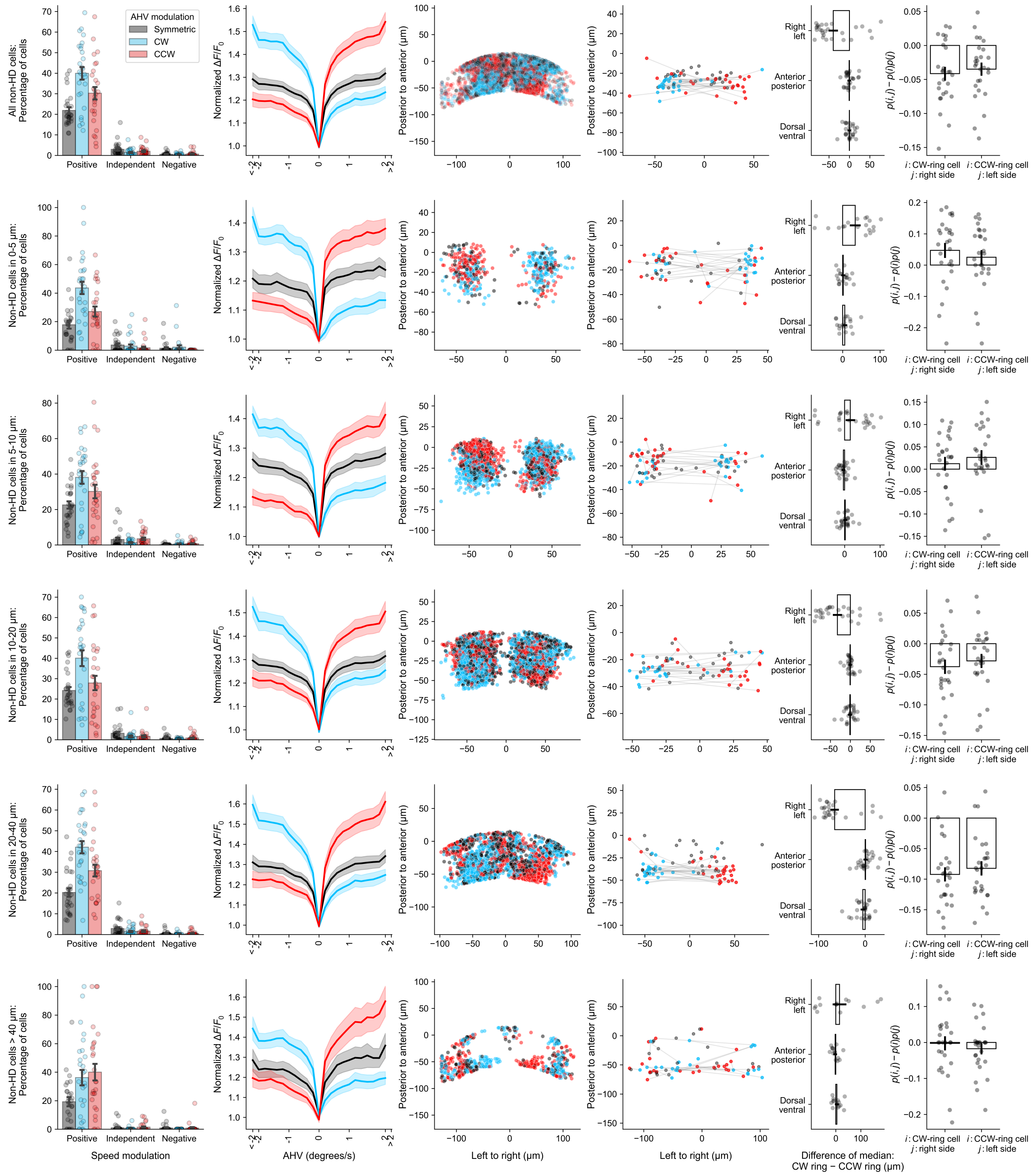

**Fig. S2** | Properties of non-HD cells. The cells are grouped according to their distances to the nearest HD cells.
